# Gene expression variation underlying tissue-specific responses to copper stress in *Drosophila melanogaster*

**DOI:** 10.1101/2023.07.12.548746

**Authors:** Elizabeth R Everman, Stuart J Macdonald

## Abstract

Copper is one of a handful of biologically necessary heavy metals that is also a common environmental pollutant. Under normal conditions, copper ions are required for many key physiological processes. However, in excess, copper quickly results in cell and tissue damage that can range in severity from temporary injury to permanent neurological damage. Because of its biological relevance, and because many conserved copper-responsive genes also respond to other non-essential heavy metal pollutants, copper resistance in *Drosophila melanogaster* is a useful model system with which to investigate the genetic control of the response to heavy metal stress. Because heavy metal toxicity has the potential to differently impact specific tissues, we genetically characterized the control of the gene expression response to copper stress in a tissue-specific manner in this study. We assessed the copper stress response in head and gut tissue of 96 inbred strains from the *Drosophila* Synthetic Population Resource (DSPR) using a combination of differential expression analysis and expression quantitative trait locus (eQTL) mapping. Differential expression analysis revealed clear patterns of tissue-specific expression, primarily driven by a more pronounced gene expression response in gut tissue. eQTL mapping of gene expression under control and copper conditions as well as for the change in gene expression following copper exposure (copper response eQTL) revealed hundreds of genes with tissue-specific local *cis-*eQTL and many distant *trans-*eQTL. eQTL associated with *MtnA*, *Mdr49*, *Mdr50*, and *Sod3* exhibited genotype by environment effects on gene expression under copper stress, illuminating several tissue- and treatment-specific patterns of gene expression control. Together, our data build a nuanced description of the roles and interactions between allelic and expression variation in copper-responsive genes, provide valuable insight into the genomic architecture of susceptibility to metal toxicity, and highlight many candidate genes for future functional characterization.

## Introduction

Many forms of stress resistance contribute to the overall health of the individual, both through immediate consequences brought on by direct exposure to the stressor, and indirectly by increasing the risk of experiencing deleterious consequences in the future. Heavy metals are one such type of stressor. Broadly, exposure to heavy metals results in oxidative stress due to the reactive state of metal ions (Stohs and Bagchi 1995; Ercal *et al*. 2001; Jomova *et al*. 2010). Acute exposure can cause gastrointestinal distress and vomiting as well as damage to intestinal lining (Taylor *et al*. 2020; Gerhardsson 2022). Chronic heavy metal exposure has been linked to neurodegenerative diseases in humans including Alzheimer’s and Parkinson’s Disease in adults (Bonilla-Ramirez *et al*. 2011; Chen *et al*. 2016b; Bagheri *et al*. 2018; Liddell and White 2018; Bisaglia and Bubacco 2020) and learning and behavioral disorders in developing individuals (Jomova *et al*. 2010; Blaurock-Busch *et al*. 2011; Hsueh *et al*. 2017; Lee *et al*. 2018). Additionally, exposure can increase morbidity associated with health conditions including multiple sclerosis and osteoporosis (Åkesson *et al*. 2006) as well as anemia (Turgut *et al*. 2007). Heavy metal exposure risk is often related to occupation (Castilla and Nealler 1978; Neuberger *et al*. 1990; World Health Organization *et al*. 1996; Khan *et al*. 2008; Karnchanawong and Limpiteeprakan 2009; He *et al*. 2015; Knoblauch *et al*. 2017), and like many diseases ((e.g., type 2 diabetes, Crohn’s disease, heart disease, IBD (http://www.genome.gov/gwasstudies) (Hindorff *et al*. 2009; Plomin *et al*. 2009)), susceptibility to metal stress is a genetically complex trait (Zhou *et al*. 2016; Everman *et al*. 2021, 2023).

Quantitative trait locus (QTL) mapping has provided a powerful means of characterizing allelic variation that contributes to lead, cadmium, copper, and zinc resistance in several model systems including worms, flies, yeast, and plants (MacNair 1983; Jones *et al*. 2006; Bálint *et al*. 2007; Courbot *et al*. 2007; Ruden *et al*. 2009; Ehrenreich *et al*. 2012; Evans *et al*. 2018; Everman *et al*. 2021). The *D. melanogaster* model system has been leveraged in several of these studies to examine the genetic architecture of metal stress response. It is an excellent model because all of the major genes that are involved in metal detoxification play similar roles in humans, allowing for the insight gained from work in *Drosophila* to have broader applications for understanding the impact of heavy metals on human health (Balamurugan *et al*. 2004; Egli *et al*. 2006a; b; Turski and Thiele 2007; Burke *et al*. 2008; Hatori and Lutsenko 2013; Calap-Quintana *et al*. 2017; Zhou *et al*. 2017). Genome-wide association mapping in flies revealed multiple SNPs that are associated with resistance to lead and cadmium toxicity (Zhou *et al*. 2016, 2017), and we previously demonstrated that resistance to copper toxicity is influenced by multiple QTL that included several conserved genes involved in copper metabolism and homeostasis (Everman *et al*. 2021). Collectively these studies have provided valuable insight into naturally occurring genetic variants that contribute to variation in resistance to several common heavy metal pollutants. However, most studies that examine the genetic control of heavy metal resistance have focused either on the whole-animal response or on single tissues. Furthermore, work over the last decade has clearly demonstrated that the majority of sites that influence trait variation are more likely to have impacts on gene regulation rather than to result in protein coding changes (Pickrell 2014; Zhang and Lupski 2015; reviewed in Boyle *et al*. 2017; Alsheikh *et al*. 2022). Therefore, an important opportunity remains to characterize the genetic control of the gene expression response to heavy metal stress and to determine the degree to which this control varies across tissues.

Transcriptomics studies have repeatedly demonstrated that exposure to heavy metal stressors increases expression of several gene families involved in metal detoxification, metabolism, and transport as well as oxidative stress response (González *et al*. 2008; Catalán *et al*. 2016; Calap-Quintana *et al*. 2017; Qu *et al*. 2018; Everman *et al*. 2021; Felmlee *et al*. 2022; Green *et al*. 2022). However, the gene expression response to metal stress is itself also subject to genetic variation in the control of the transcriptional response, i.e., different genotypes can vary in their response to the same exposure (Findley *et al*. 2021). The loci that contribute to variation in the gene expression response can be identified using expression QTL (eQTL) mapping (Jansen 2001; De Koning and Haley 2005; Gilad *et al*. 2008). By combining RNA sequencing and eQTL mapping, we examined the tissue-specific genetic control of the gene expression response to the common metal pollutant copper using the *D. melanogaster* model system.

Although a broad oxidative stress response is expected across tissues in response to metal stress, copper is one of several heavy metals that have the potential to elicit different gene expression responses in neurological and gut tissue. This expected tissue-specific difference is in part due to the spatial distribution of specialized copper accumulation cells found in the gut of most animals including flies and humans (Calap-Quintana *et al*. 2017; Miguel-Aliaga *et al*. 2018). Genes that are responsible for maintaining normal homeostasis of the set of essential heavy metals (e.g., copper and zinc), and that play a role in the expulsion of toxic metals such as cadmium and lead, are primarily expressed in these specialized cells where they play an important protective role against excess metal ions (Calap-Quintana *et al*. 2017). Here we compared the gut response with that of head tissue due to the link between metal exposure and neurological disease in humans (Chen *et al*. 2016b), and for the opportunity to characterize differences in the tissue-specific response to oxidative stress using copper as a model metal.

Our study leverages the *Drosophila* Synthetic Population Resource (DSPR (King *et al*. 2012a)), a panel of multiparental, advanced intercross strains exhibiting dramatic variation in copper resistance that we have previously employed to map numerous QTL for resistance (Everman *et al*. 2021). Here we build on this study and use a subset of 96 strains to characterize the regulatory control of copper stress. Since the DSPR is composed of genetically stable inbred strains, it is especially well suited to eQTL mapping studies that quantify expression across tissues and treatments. We identified thousands of eQTL in both gut and head tissue using animals exposed to either control or copper-exposure treatments. Tissue-specific genetic control of gene expression was extensive, suggesting that regulatory variation has the potential to have profound, yet context-specific effects on susceptibility to heavy metal stress. Ultimately, our characterization of the genetic control of gene expression response to copper stress provides nuanced insight into the genetic control of heavy metal resistance and its potential health consequences.

## Methods

### Fly stocks

We used the *Drosophila* Synthetic Population Resource (DSPR), a large multiparental mapping panel with >1500 recombinant inbred lines (RILs) (see King *et al*. 2012a, b for additional details). In a previous study, we exposed 60 adult females from each of 1556 RILs (767 and 789 RILs from the A and B populations, respectively) to 50mM CuSO_4_, expressing resistance as the percentage of adults alive at 48 hours (Everman *et al*. 2021). For the present study, we selected 48 of the “resistant” RILs from the B population that were in the top 25% of the distribution of adult copper resistance and 48 of the “sensitive” RILs from the bottom 25% of the distribution (Figure S1).

### Rearing and assay conditions

Flies were reared and assayed across multiple batches in the same incubator at 25°C and 50% humidity with a 12:12hr light:dark photoperiod. To maintain consistency with prior work, experimental females for the present study were obtained from each of the 96 DSPR RILs in the same manner used to measure adult copper resistance (Everman *et al*. 2021). Briefly, adults were placed on cornmeal-molasses-yeast media and were allowed to oviposit for 2 days before being discarded. Experimental females from the following generation were sorted over CO_2_ into groups of 20 and allowed to recover for 24 hours on fresh cornmeal-molasses-yeast media. Following recovery, 3-5 day old females were transferred to Instant *Drosophila* Media (Carolina Biological Supply Company 173200) hydrated with either water (control) or 50mM CuSO_4_ (Copper(II) sulfate, Sigma-Aldrich C1297) and exposed to treatment conditions for 8 hours, after which tissues were harvested (see below). The 8-hour exposure period lasted from lights on (0800hrs) to 4 hours before lights off. No flies died during the 8-hour exposure period.

Strains were assayed in a series of small batches to accommodate the time required for tissue processing following the exposure period, with sensitive and resistant RILs evenly distributed across each batch. Given the different processing techniques for heads and gut, we obtained these tissues from different individuals of the same genotypes in different batches. The average batch size for head collection was 25 DSPR RILs and average batch size for gut dissection was 6 DSPR strains.

### Tissue collection, RNA isolation, and sequencing library preparation

To obtain head tissue, we exposed 60 females per DSPR RIL to control and copper conditions (three vials of 20 females per treatment) for 8 hours. At the end of the exposure period, flies from each treatment and RIL were moved to a labeled screw-top 50mL tube, flash frozen in liquid nitrogen, immediately vortexed to separate heads from bodies, and then stored at -80°C for up to five days. We used a series of three-inch diameter stacking brass sieves – chilled on dry ice for at least five minutes to prevent tissue thawing – to isolate the heads from each sample (see Supplementary Methods). Heads for each RIL/treatment were dispensed into a screw-top microcentrifuge tube containing 3-4 glass beads and held on dry ice. Subsequently, we added 300μL TRIzol Reagent (Invitrogen, 15596018) to each sample and stored samples at -80°C until RNA extraction.

To obtain gut tissue, we exposed 20 females per DSPR RIL to control and copper conditions for 8 hours. Guts, including the fore-, mid-, and hindgut, were dissected from 10 live individuals per RIL/treatment combination in 1X PBS and placed in screw-top microcentrifuge tubes containing 3-4 glass beads and 300μL TRIzol. Samples were chilled on ice during dissection and stored at -80°C until RNA extraction.

RNA was extracted from each of the 384 samples in batches of 30 samples using the Direct-zol RNA Miniprep kit (Zymo Research, R2050). Batches consisted of samples from the same tissue and were processed in the order individual samples were collected. RNA was eluted in 50μL water, and concentrations were determined via a NanoDrop spectrophotometer and then standardized to 20ng/μL in 96-well plates. Because head and gut tissues were not collected contemporaneously, two plates contained samples from heads and two contained samples from gut tissue. The order of the samples from each RIL and treatment was largely consistent across the head and gut sets of plates; all plates included both copper and control samples for a given RIL and held an even representation of sensitive and resistant RILs (Table S1).

Half-reaction, unique dual indexed TruSeq Stranded mRNA sequencing libraries (Illumina, 20020595) were generated and sequenced by the University of Kansas Genome Sequencing Core. Concentrations for every library were obtained using the Qubit dsDNA HS assay kit (ThermoFisher, Q32854), and successful libraries were pooled based on concentration within each of the four tissue-specific plates described above (Table S1). This resulted in two pools of head libraries (one 94- and one 96-plex), and two pools of gut libraries (one 93- and one 96-plex). Paired-end 37bp reads were obtained by sequencing each pool separately on NextSeq550 High-Output flowcells. Raw read counts for the 94-plex Head, 93-plex Gut, and 96-plex Gut libraries were 5.1 – 5.6 million PE37 read pairs per sample (Figure S2). The initial sequencing run of the 96-plex Head library pool under-clustered, so the pool was sequenced a second time on a NextSeq550 Mid-Output flowcell. Assessment of the reads revealed no difference in our ability to map reads from either run (Figure S3), so we combined reads across the two runs, resulting in an average of 3.8 million PE37 read pairs per sample for this library pool (Figure S2).

### Sequencing data preprocessing

#### Read alignment and gene filtering

Quality assessment, adapter removal and read trimming (retaining reads with a minimum of 15 bp, Figure S4) were performed with fastp (v. 0.20.1) (Chen *et al*. 2018). Only paired reads were retained. Filtered read pairs were aligned in a variant-aware manner to the *Drosophila* reference genome (Release 6.33) using HISAT2 (v. 2.1.0) (Kim *et al*. 2019). SNP variants identified in the DSPR founders (http://wfitch.bio.uci.edu/~tdlong/SantaCruzTracks/DSPR_R6/dm6/variation/DSPR.r6.SNPs.vcf.gz) were included in the HISAT2 genome index following instructions provided with the software. Aligned reads were sorted with SAMtools (v. 1.9) (Li *et al*. 2009) and quantified with featureCounts (v. 2.0.1) (Liao *et al*. 2014). Post alignment and quantification analyses were all performed in R (v. 4.1.3) (R Core Team 2017). Average library size prior to additional filtering was 4.03 million reads. In R, genes with low expression were filtered out by first removing genes with zero counts across all samples and then by removing genes with fewer than 10 counts in 47 samples (the number of samples unique to each of the three-way interaction categories, e.g. low copper resistance head tissue samples exposed to copper stress (Chen *et al*. 2016a)). Following filtering, 9842 genes were retained for downstream analyses.

#### Comparison between alignment pipelines

We also aligned fastp-filtered reads using the kallisto (v. 0.46.2 (Bray *et al*. 2016)) pseudoalignment pipeline and the Ensembl *Drosophila* transcriptome release 90. Average library size was 3.96 million reads following kallisto alignment. We applied the same post-alignment filtering parameters described above to the kallisto generated counts. Because kallisto alignment generates output quantified by transcript, filtering resulted in retention of a slightly different set of genes totaling 9913 genes, 9434 of which were shared with the filtered HISAT2 gene set. To compare gene expression results from the HISAT2 and kallisto pipelines, we calculated gene-wise correlations in expression following log2 transformation and quantile normalization across the 379 libraries. Sample and tissue-specific read counts for the 9434 genes retained in both the kallisto and HISAT2/SAMtools/featureCounts pipeline were highly correlated (mean R = 98.2%, Figure S5). Correlations for normalized gene counts within each tissue/resistance class/treatment set of samples were also high (96% - 97%). The 267 genes with counts that were less than 90% correlated between the two pipelines were not enriched for any functional or molecular categories following gene ontology analysis (Flymine, (Lyne *et al*. 2007)), suggesting that differences in gene expression counts were not due to systematic discrepancies between pipelines. Given the high consistency between pipelines, all downstream differential expression (DE) analyses and eQTL mapping presented below are derived from the HISAT2/SAMtools/featureCounts pipeline.

### Effect of tissue, treatment, and resistance class on gene expression

Gene expression was normalized using the weighted trimmed mean of M-values (mean of log expression ratios (Robinson and Oshlack 2010)), and DE analysis was performed with limma (Ritchie *et al*. 2015) by fitting the full-factorial linear model (Expression ∼ Tissue * Treatment * Resistance class + Sample Pooling). The “Sample Pooling” model term accounts for technical variation due to plate, library pool and/or sequencing flowcell (these 3 factors cannot be distinguished in our design). Genes with expression variation significantly influenced by each term in the model were identified by fitting contrasts to the full model in limma with the contrasts.fit function. This step was followed by the eBayes function, which uses an empirical Bayes method to calculate log fold change in normalized gene expression and to determine significance between each group identified by the contrast (contrast order: expression in gut relative to head tissue, copper relative to control treated samples, sensitive relative to resistant strains). Significant DE was determined using adjusted P values to account for multiple comparisons (Benjamini-Hochberg multiple test correction, FDR threshold = 5%).

### Tissue and treatment-specific eQTL analysis

We performed eQTL mapping for 6 separate datasets, testing different numbers of gene expression traits in each due to some genes having tissue- and/or treatment-specific expression (Head-Control: 9783, Head-Copper: 9842, Head-Response: 9830, Gut-Control: 9842, Gut-Copper: 9842, Gut-Response: 9841 genes). The “Response” to copper treatment was calculated as gene expression under copper conditions minus expression under control conditions for all paired samples. Positive Response values indicate genes with higher expression under copper conditions (copper treatment-induced), while negative Response values indicate genes that are repressed by copper treatment. Each of the six datasets was separately quantile normalized following log2 transformation. To preserve the direction of gene expression change in both Response datasets, we performed log2 transformation and quantile normalization on the absolute values of the copper-response expression and reassigned the sign following normalization.

We corrected expression measures for variation due to technical and environmental factors using principal components analysis (PCA; described in further detail below). Separately for each dataset, gene expression was corrected by regressing out the effects of PCs that explained more than 2% of the variance in gene expression, or that were correlated with known technical factors. eQTL mapping was performed on quantile normalized residual gene expression for each of the six datasets using R/qtl2 (Broman *et al*. 2019). eQTL mapping treats each gene expression measure as a trait, and we test for associations between genotype and variation in gene expression at each genome marker (every 10kb in the DSPR (King *et al*. 2012a)). For each gene, we regressed expression on additive probabilities indicating the likelihood that a particular region of the genome was inherited from one of the 8 DSPR founders. Gene-specific significance thresholds were assigned by permuting gene expression estimates among the DSPR strains 1000 times.

Given the modest sample size for each eQTL analysis (93-96 DSPR strains), we defined QTL intervals using a 3-LOD drop, since a 2-LOD drop can give overly narrow intervals when fewer RILs are assayed (King *et al*. 2012a). *cis* eQTL were defined as above-threshold peaks for which the upper or lower boundary of the 3-LOD drop was within 1.5cM of the target gene, while peaks outside this interval were classified as distant or *trans* eQTL. We note that broader peak intervals result in more peaks being designated as *cis* eQTL and fewer peaks as *trans* eQTL (King *et al*. 2014; King and Long 2017). Peaks were removed if they consisted of a single, above-threshold marker, or if the peak position was outside the lower and upper peak boundary (such phenomena are more common in experiments with limited sample size and are often found near telomere and centromere regions where the impact of low power is exacerbated). eQTL peaks, estimated founder effects, and percent variance were identified and calculated using custom code derived from the DSPRqtl (King *et al*. 2012b) and R/qtl2 mapping programs.

We examined the six datasets for evidence of eQTL that were shared among datasets. Comparisons were made between tissues within treatment (Head-Control vs Gut-Control, Head- Copper vs. Gut-Copper, Head-Response vs. Gut-Response) and within tissue between treatments (Heads-Control vs Heads-Copper and Gut-Control vs Gut-Copper). Our ability to identify shared eQTL across multiple datasets is influenced by power to detect eQTL and the effect size of the eQTL. In an attempt to account for increased uncertainty, eQTL were considered shared if the peak positions were within 1.5cM of each other and/or if the peak intervals overlapped.

### PCA correction of gene expression for eQTL analysis

A common strategy to reduce spurious gene-eQTL associations in high-dimensional data is to use PCA to account for known and unknown technical or environmental factors (Leek and Storey 2007; Pickrell *et al*. 2010; Gaffney *et al*. 2012; King *et al*. 2014). To determine the most appropriate PCA correction for our data, we compared three versions of the eQTL mapping analysis using 1) uncorrected data, 2) data corrected for PCs that explained more than 2% of the variance and/or were correlated with tissue collection batch or sample pool (general linear model test of the effect of technical variation on PC, uncorrected α = 0.05) and 3) data corrected for PCs that explained more than 1% of variance and/or were correlated with technical variation. The set of PCs accounted for in each case are listed in Table S2. For brevity, we will simply refer to the two PCA-corrected versions of the dataset as 2% and 1% corrected. Corrections were made by regressing out PCs separately for each of the six expression datasets. Residuals were quantile normalized once more and treated as gene expression values for each of the eQTL analyses and for all downstream analyses.

We performed the three eQTL analyses (no correction, 2% correction, 1% correction) for each of the six datasets as described above. In Head-Control, Head-Copper, Gut-Control, and Gut- Copper eQTL analyses, failing to correct for technical variation via PCA resulted in many more peaks far from the target gene compared to the 2% and 1% cutoffs. Because not correcting for technical factors increases noise, it is likely the large number of distant peaks in the uncorrected data includes many more false positives than true *trans* eQTL (Figure S6). The relationship between peak number and data correction strategy was more pronounced for head samples, likely due to the known technical effect of sequencing pool (Figure S6) that we account for in the PCA-corrected versions. Head- and Gut-Response data were less sensitive to the PCA correction (Figure S6, S7). Removing PCs explaining 1% of the variance further reduced the total number of peaks in both head and gut analyses relative to the 2% variance correction (Figure S7). However, this difference was slight, and overall levels of noise observed in individual LOD plots were visually and statistically similar between the 2% and 1% data correction (average correlation in genome-wide LOD scores between 1% and 2% corrected data for genes with eQTL ranged from 70% - 83%; Figure S8). To avoid removing true positives through use of a strict 1% cutoff, all subsequent analyses/results employ the 2% PCA-corrected version of the dataset.

### Identification of eQTL hotspots

Identification of eQTL hotspots in each of the six eQTL mapping datasets entailed two initial steps. First, we used a chi-square goodness of fit test, assessing each dataset separately, to determine if *trans* eQTL were overrepresented in any of the 2cM intervals distributed along each chromosome arm. Second, because the chi-square test only indicates whether any interval on the chromosome arm had an observed number of eQTL that deviated from expected number of eQTL, we calculated the percent contribution of each interval to the overall chi-square value. Percent contribution was calculated for each interval as

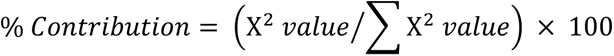

Intervals with a percent contribution value greater than 10% were considered potential hotspots. We chose 10% as a permissive threshold for detecting hotspots based on a visual assessment of percent contribution values as they varied across the chromosome arm. In most cases, our approach highlighted one interval per chromosome arm with enrichment for *trans* eQTL.

We took a three-step approach to further characterize *trans* eQTL hotspots and to identify potential regulatory candidate genes responsible for hotspots. Each step was carried out independently for each hotspot. First, we created a composite variable using PCA to summarize expression variation for all genes with a *trans* eQTL in that hotspot. We then used QTL mapping in the same manner as described above to isolate QTL for this composite variable (PC1), and if a peak was detected that overlapped the location of the hotspot, the hotspot was retained in subsequent steps.

Second, we repeated QTL mapping of the composite variable, but now iteratively tested whether each gene that fell within the hotspot interval had a mediating effect on the composite QTL peak by including the expression level of each gene as a covariate in the QTL analysis. If adding a gene expression covariate eliminated the composite variable QTL (or dramatically reduces the LOD score at the peak), the gene was a candidate mediating gene of the *trans* eQTL hotspot. The degree to which the LOD score was reduced by potential mediating genes varied across all the hotspots, making it difficult to establish a consistent cutoff value. We therefore assessed sets of mediating genes separately for each hotspot, and identified candidates that had the most pronounced mediating effect at the composite peak position. If several genes had similar mediating effects on the composite peak, we included them as candidates. This permissive approach produced a list of genes that should be considered for future follow up studies to further distinguish between true mediating genes versus those that have correlated expression with mediating genes. Because the 2cM intervals used to initially identify hotspots were created arbitrarily, we used the boundaries of the *trans* eQTL peak intervals to define the genomic region associated with the hotspot by using the most minimum and most maximum interval boundary for all *trans* eQTL in each hotspot. All the genes that fell within these expanded regions were included in this second step except for those genes that did not appear in our gene expression dataset. Some genes with very low levels of expression were removed from our expression datasets during filtering and so were not included in this analysis (see above). Covariate data used in each analysis corresponded to the dataset in which the hotspot was identified. For example, gene expression in gut tissue under copper conditions was used as covariate data to account for a composite variable derived for a hotspot detected in the Gut- Copper dataset.

Third, we further examined potential mediating genes by testing the correlation between estimated founder haplotype effects at any *cis* eQTL that happened to be associated with the potential mediating gene that fell within the hotspot interval. As in the second step, we used the *cis* eQTL that corresponded to the dataset in which the *trans* eQTL hotspot was detected.

### Gene annotations and ontology analyses

We obtained annotation and ontology information for focal sets of genes highlighted by differential expression analysis, eQTL mapping, and assessment of eQTL hotspots using the *Drosophila melanogaster* annotation tool available from biomaRt (v. 2.50.3 (Durinck *et al*. 2005)) via Ensembl (Cunningham *et al*. 2022) and the orb.DM.eg.db R package (v. 3.14.0 (Carlson 2019)). Gene enrichment analyses were performed using the R package GOstats (v. 2.60.0 (Falcon and Gentleman 2007)), which uses hypergeometric tests for overrepresentation of GO terms with a correction for multiple tests.

### Data availability

RNAseq reads are available from NCBI SRA BioProject PRJNA993605. All data including DE results, quantile-normalized expression data used for eQTL analyses, and code used to perform eQTL analyses are available from FigShare. Supplementary material contains Supplementary Methods that provide more detail on the isolation of head tissue as well as the alignment pipeline, code to carry out eQTL mapping analysis, and all input and output data generated for each analytical step.

## Results

### Effect of tissue, treatment, and resistance class on gene expression

A principal goal of our study was to characterize the gut- and head-specific gene expression responses to copper exposure in genetically diverse copper-sensitive and copper-resistant DSPR strains. Our differential expression model tested the three-way interaction between tissue (gut, head), treatment (control, 8-hour exposure to 50mM CuSO_4_), and DSPR strain resistance class (resistant, sensitive). The primary model parameters that influenced expression were tissue and treatment. Gene expression was strikingly distinct between gut and head samples resulting in differential expression (DE) of 8915 of the 9842 genes tested (91% of genes, 5% FDR; Figure 1A). Treatment alone influenced gene expression in 1156 genes (12%, Figure 1B), while resistance class alone influenced expression of three genes: *asRNA:CR44107*, *CG18563* (involved in serine-type endopeptidase activity (Thurmond *et al*. 2019)), and *CG6023*.

**Figure 1.**
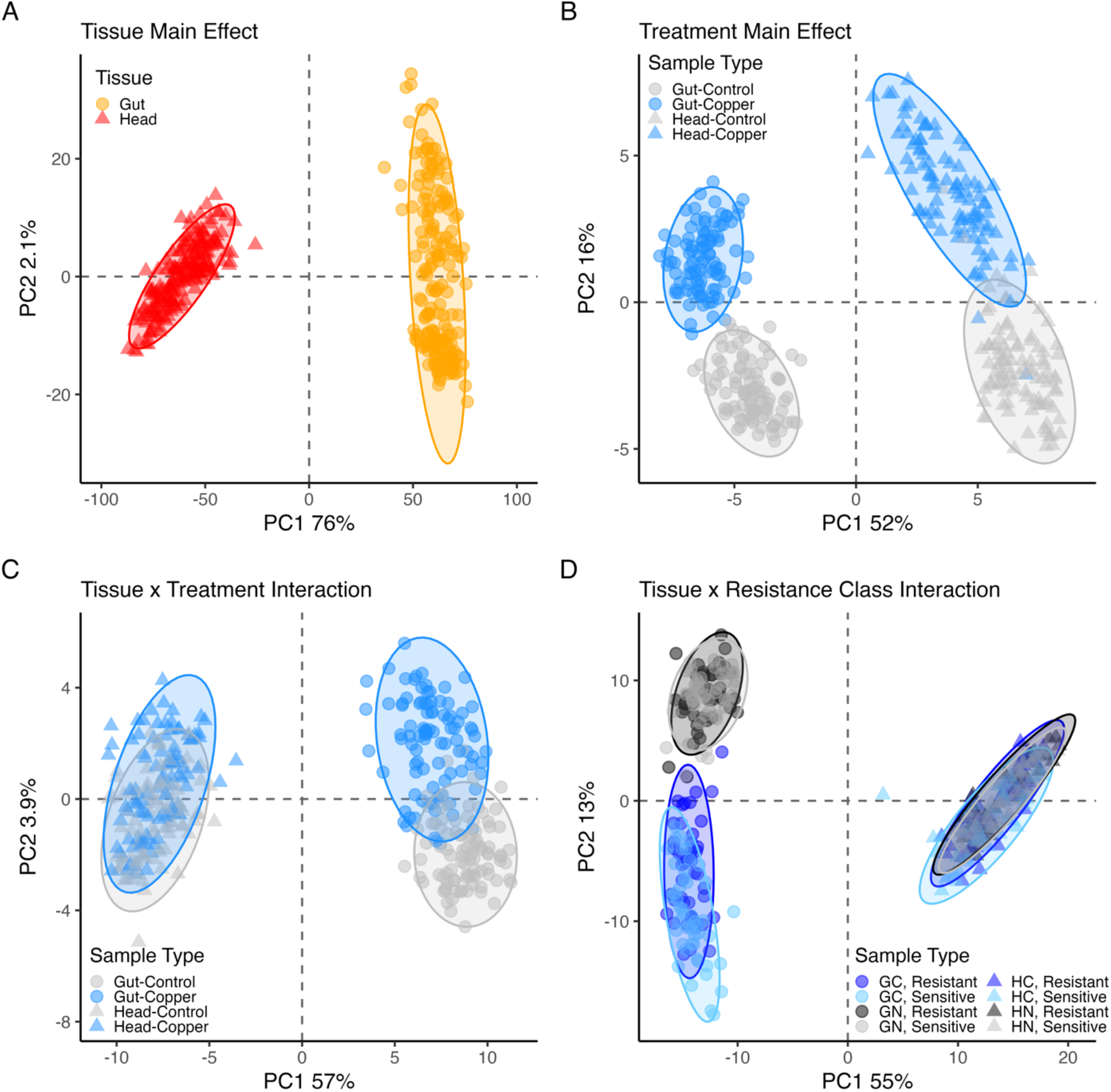
Gene expression varied between tissues and treatments. Here we employ PCA to depict variation in gene expression in the top differentially-expressed genes with FDR < 0.05 and log fold change > 1 for four significant model terms. Each PCA was carried out with a different set of genes: A. 5031 genes, B. 59 genes, C. 113 genes, D. 341 genes. The effect of tissue on gene expression is shown in (A) but is also evident in all other plots. B. Gene expression shifted in both head and gut tissue in response to copper exposure. C. Gene expression shifted in response to copper treatment in both head and gut tissue. Shifts were typically in the same direction in both tissues with a slightly higher magnitude response observed for genes in gut tissue relative to head, leading to the interaction. D. Tissue and resistance class interacted to influence gene expression, although the effect of resistance class was slight. The significant interaction was driven by a larger magnitude difference in expression between sensitive and resistant strains in the gut tissue in response to copper. In all plots, triangle symbols indicate head samples, circles indicate gut samples. Log2, quantile-normalized gene expression data were corrected for the effect of sequencing pool prior to clustering analysis.

We also found that for 70% of genes, expression was influenced by an interaction between tissue and treatment (Figure 1C). On average, this tissue x treatment interaction was driven by a response to copper exposure that was in the same direction in head and gut tissue, but that was more pronounced in gut tissue (Figure 1C). This pattern is consistent with the slightly larger effect of treatment alone in gut tissue (Figure 1B) and might be expected because flies were exposed to copper via ingestion in our assay. The effect of resistance class varied subtly between gut and head tissue, influencing 32% of genes (Figure 1D), with the interaction driven by a slightly greater effect of resistance class in gut tissue versus head tissue. The treatment and resistance class interaction did not influence the expression of any gene tested, and the three-way interaction between tissue, treatment, and resistance class influenced 13 genes (0.13%), of which 8 had known or predicted functions. None of the genes influenced by the three-way interaction were known copper response genes or had clear connections to metal toxicity response. Information on all DE genes is available in the Supplementary data.

### Differential expression of copper-related genes

Seven genes that have been previously linked to binding, metabolism, and detoxification of copper were among the DE genes influenced by treatment (Figure 2A) and 28 genes in these categories were influenced by the tissue by treatment interaction (Figure 2B). For several genes that have been previously investigated in the context of copper toxicity, the change in expression across treatments/tissues was in the expected direction. For instance, *Syx5* is a critical copper homeostasis gene that is required for proper accumulation of copper ions under normal (copper- scarce) conditions and is hypothesized to aid in proper localization of copper import proteins (Norgate *et al*. 2010). Norgate *et al*. (2010) demonstrated reduced expression of *Syx5* increased resistance to excess copper. Complementing this previous work, we found exposure to copper stress significantly reduced expression of *Syx5* in both head and gut tissues (Figure 2A), suggesting that downregulation of *Syx5* in the whole organism may result from copper stress.

**Figure 2.**
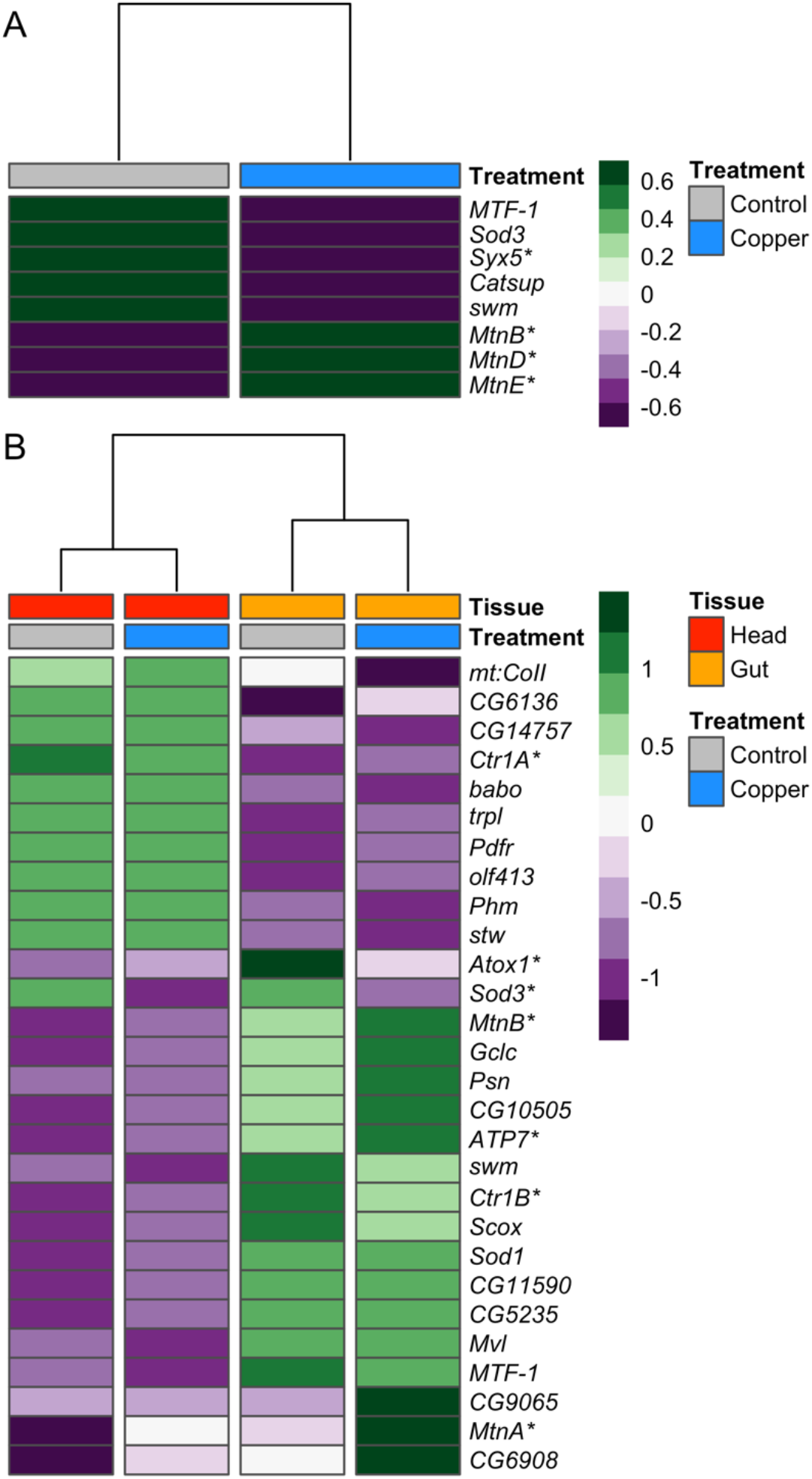
Treatment and the interaction between treatment and tissue influenced expression of several genes previously shown to be involved in detoxification, homeostasis, or binding of copper and other heavy metals. A. Heatmap of genes differentially expressed due to treatment that have been previously associated with copper ion response. B. Heatmap of genes differentially expressed due to the interaction between tissue and treatment. In both heatmaps, tissue and treatment groups are indicated at the top of the heatmaps, and gene expression is presented as average normalized expression for strains belonging to the two resistance classes. Asterisks beside gene names highlight genes discussed in text.

Expression of the metallothionein family genes in response to copper stress was also consistent with expectations. Copper exposure was expected to increase expression of the metallothionein genes, which are involved in the sequestration of excess copper ions as a first defense against copper toxicity (Egli *et al*. 2006a; Calap-Quintana *et al*. 2017). Further, expression was expected to be higher in gut compared to head tissue because primary expression of metallothioneins has been shown to occur in specialized copper accumulating cells that line the midgut in flies (Calap-Quintana *et al*. 2017) As expected, we found that expression of four metallothioneins (*MtnA*, *MtnB*, *MtnD*, and *MtnE*) was significantly increased in response to copper exposure in both head and gut tissue but the increase in expression was more pronounced in gut tissue (Figure 2).

Three key copper transporters (*Ctr1A*, *Ctr1B*, and *ATP7*) were also among the genes influenced by an interaction between tissue and treatment that followed expected expression patterns based on previous reports. The ATP7 and Ctr1B copper transporters function primarily in specialized copper-accumulating cells that line the intestine in mammals and the midgut in flies (Zhou *et al*. 2003; Turski and Thiele 2007; Calap-Quintana *et al*. 2017), whereas the Ctr1A copper transporter is ubiquitously expressed in both mammals and flies (Turski and Thiele 2007; Calap-Quintana *et al*. 2017). Under control and copper conditions, we observed higher expression of *ATP7* and *Ctr1B* in gut tissue relative to head tissue and higher expression of *Ctr1A* in head tissue compared to gut tissue (Figure 2B). Our data are also consistent with previous reports that *Ctr1A/B* copper importers are downregulated in *D. melanogaster* in response to copper overexposure (Zhou *et al*. 2003; Calap-Quintana *et al*. 2017) as we observed a decrease in expression of both *Ctr1A* and *Ctr1B* following copper exposure. The pattern of decreased expression in response to copper was consistent across tissue but was more pronounced in head tissue (*Ctr1A*) or gut tissue (*Ctr1B*) depending on the gene. Although *ATP7* expression under control conditions followed tissue-specific expectations, we observed that exposure to copper stress significantly increased *ATP7* expression, a pattern that was more pronounced in gut tissue (Figure 2B). *ATP7* is a major copper exporter (Zhou *et al*. 2003), facilitating the transfer of copper from copper accumulating cells in the gut to other tissues. Increased copper export under stressful conditions may play an important role in the response to copper toxicity.

Expression of the copper chaperone *Atox1* was also differentially affected by treatment and tissue. As proteins, Atox1 transports copper to ATP7 (Calap-Quintana *et al*. 2017; Kamiya *et al*. 2018) and to the extracellular antioxidant enzyme SOD3 (Itoh *et al*. 2009; Kamiya *et al*. 2018). Both *Atox1* and *Sod3* were more highly expressed in gut tissue than in head tissue (Figure 2B), and exposure to copper stress led to a decrease in expression of both genes. Work presented by Itoh *et al*. (2009) suggests that this shift in expression may be mechanistically linked because Atox1 functions both to transfer copper to SOD3 and to regulate its expression as a copper-dependent transcription factor (Itoh *et al*. 2009). Under normal conditions, null mutations in *Atox1* have been shown to lead to copper deficiency, and excess copper led to decreased Atox1 protein expression in wild-type flies (Hua *et al*. 2011).

To characterize the tissue-specific stress responses more broadly we used gene ontology analysis of those genes showing significant tissue by treatment interactions in our differential expression analysis. Of the 6362 genes with tissue- and treatment-specific patterns in expression, 778 could be categorized into one or more stress response categories (Table 1). As expected, when examined on a gene-by-gene basis within each category we often saw different responses to copper stress in head and gut tissue, indicating a range of tissue-by-treatment interactions (Figure 3A). However, when the average response of genes in each GO category was considered, we found that the overall expression response to copper treatment was often more pronounced in gut tissue compared to head tissue (Figure 3B). This pattern was observed for regulation of and response to stress (both cellular and general), response to endoplasmic reticulum stress, and categories related to stress-activation of the MAPK and protein kinase signaling cascades (Figure 3B). In contrast, copper-induced shifts in expression of genes involved in the response to oxidative stress was more pronounced in head tissue compared to gut tissue (Figure 3B). Importantly, each of the oxidative stress response genes also contribute to each of the categories that showed a more pronounced response in gut tissue. This implies that oxidative stress response is also elevated in gut tissue, but it is one stress response pathway among many that are activated in the gut, whereas in head tissue oxidative stress response is singularly pronounced.

**Figure 3.**
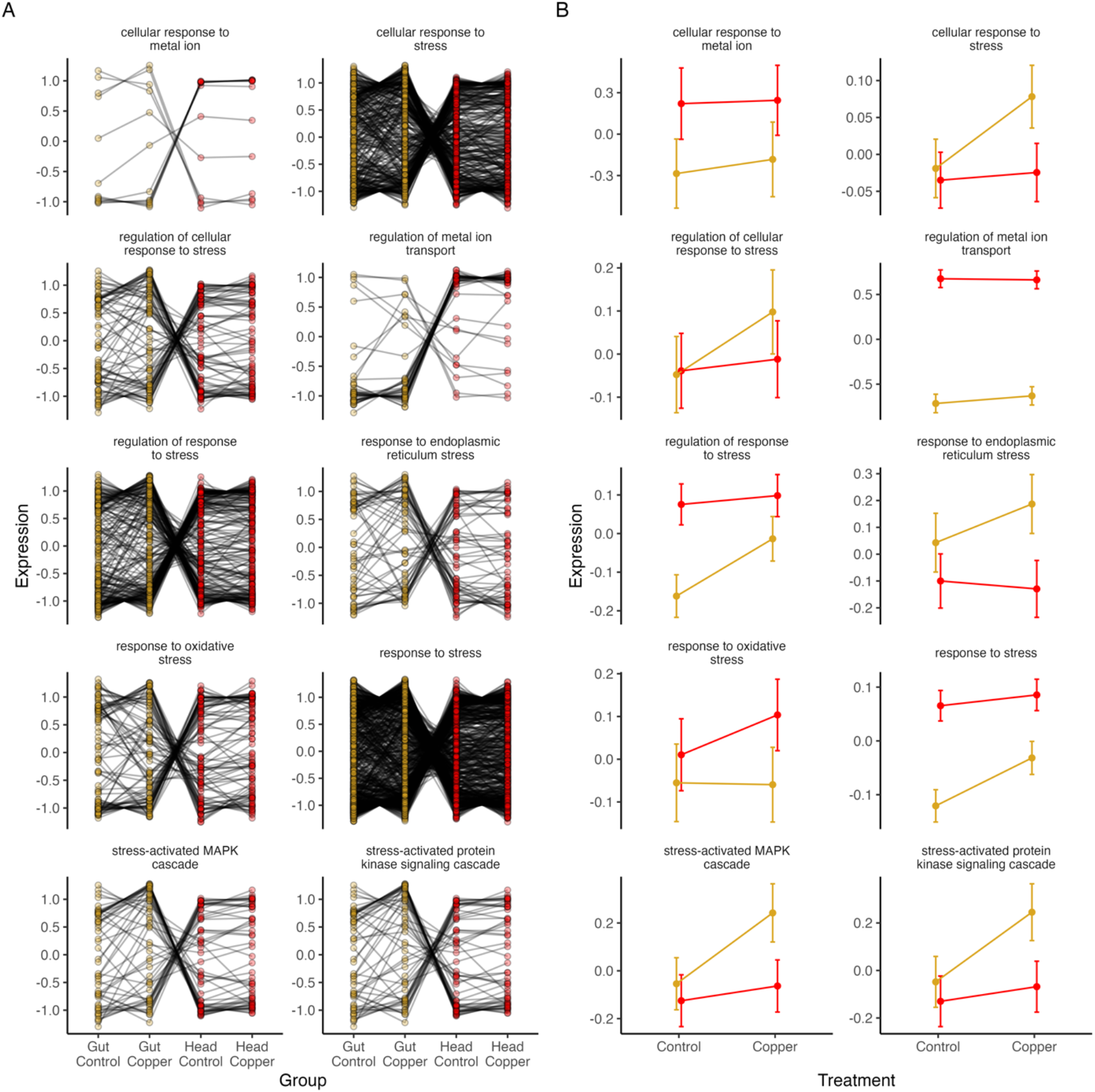
778 genes with a significant tissue by treatment interaction effect could be classified by their involvement in stress response pathways. A. At the gene level, mean expression across DSPR strains under copper and control conditions in gut and head tissue demonstrated many distinct tissue and treatment specific patterns of expression response to copper. B. When expression was summarized at the level of GO category, we found that most genes that are involved in cellular response to stress, regulation of stress response, response to endoplasmic reticulum stress, and involvement in stress-activated MAPK and protein kinase signaling had a more pronounced response in gut tissue compared to head tissue. Conversely, genes involved in the response to oxidative stress had a more consistently pronounced response in head tissue compared to gut tissue. Data shown are mean expression response for all genes in each treatment and tissue combination +/- SE. In both plots, gold indicates gut samples; red indicates head samples.

**Table 1.** Enrichment of stress-related GO categories (5% FDR) in genes with a tissue by treatment interaction.

| Term | GO ID | N Genes in<br>GO Category | N Genes (%)<br>with sig<br>Interaction | P Value |
| --- | --- | --- | --- | --- |
| Response to stress | GO:0006950 | 1321 | 735 (56%) | $<2.47e^{-24}$ |
| Regulation of response to stress | GO:0080134 | 346 | 218 (63%) | $<5.97e^{-15}$ |
| Cellular response to stress | GO:0033554 | 646 | 370 (57%) | $<6.83e^{-15}$ |
| Response to oxidative stress | GO:0006979 | 140 | 91 (65%) | $<5.83e^{-8}$ |
| Regulation of metal ion transport | GO:0010959 | 57 | 40 (70%) | $<2.16e^{-5}$ |
| Response to endoplasmic reticulum stress | GO:0034976 | 89 | 57 (64%) | $<3.27e^{-5}$ |
| Stress-activated protein kinase signaling cascade | GO:0031098 | 88 | 56 (64%) | $<4.99e^{-5}$ |
| Regulation of cellular response to stress | GO:0080135 | 131 | 78 (60%) | $<5.95e^{-5}$ |
| Stress-activated MAPK cascade | GO:0051403 | 87 | 55 (63%) | $<7.58e^{-5}$ |
| Cellular response to metal ion | GO:0071248 | 15 | 13 (87%) | $<5.68e^{-4}$ |

Overall, we found pervasive differences in expression between tissues, with many genes – including candidate metal responsive genes – showing an expression change in response to copper treatment as well as variation in copper response among tissues. We previously reported a minor effect of DSPR strain-specific copper resistance on the gene expression response to copper (Everman *et al*. 2021). Classifying DSPR strains into copper resistant and susceptible classes in the present study explained relatively little of the expression variation, and resistance class influenced only a small number of genes, none of which are members of known metal response pathways. Many of the key copper response genes highlighted above and in Figure 2 and Figure 3 have essentially equal expression in resistant and sensitive strains, suggesting that the base level toxicity response to our copper treatment is similar between resistant and sensitive strains assayed in this study.

### Expression QTL mapping

#### Properties of mapped eQTL

eQTL mapping was executed for 6 datasets: Head-Copper, Head-Control, Head-Response, Gut-Copper, Gut-Control, Gut-Response (see Methods for additional detail). In total over all datasets, 4377 genes were associated with at least one eQTL and 98% of these 4377 genes were differentially expressed due to one or more of the DE model terms and interactions (described above). For most genes (83-98%; Table 2), we detected one eQTL (either *cis* or *trans*) per gene per dataset (Figure S9). Genes with more than one eQTL per dataset included a multidrug resistance gene *Mdr65* and two cytochrome p450 genes *Cyp6t3* and *Cyp12d1-d*, which have been linked to response to insecticides (Daborn *et al*. 2007; Esteves *et al*. 2021). Two genes linked to copper ion binding and homeostasis (*Mco1* and *CG6908* (Lang *et al*. 2012; Thurmond *et al*. 2019)) were also among the genes with multiple eQTL per dataset. However, no gene enrichment related to metal detoxification or response was evident in genes with more than one eQTL peak per dataset. Gene annotation and GO analysis for genes with multiple eQTL is available in the Supplementary data.

**Table 2.** Summary statistics for eQTL mapping analyses.

| Analysis | eQTL Type | N eQTL Peaks | N genes with eQTL | Total N unique genes with eQTL | % Genes with 1 eQTL peak | Mean Percent Variance (range) |
| --- | --- | --- | --- | --- | --- | --- |
| Gut-Control | *cis* | 1784 | 1779 | 1993 | 96% | 43.1 (25.0-74.4) |
| Gut-Control | *trans* | 295 | 286 |  |  | 33.7 (10.4-63.0) |
| Gut-Copper | *cis* | 1677 | 1674 | 1860 | 97% | 42.4 (22.0-76.0) |
| Gut-Copper | *trans* | 248 | 237 |  |  | 33.4 (21.2-68.2) |
| Gut-Response | *cis* | 139 | 138 | 316 | 98% | 40.0 (25.5-72.5) |
| Gut-Response | *trans* | 183 | 180 |  |  | 32.3 (16.6-41.9) |
| Head-Control | *cis* | 1875 | 1869 | 2125 | 96% | 43.6 (22.8-76.8) |
| Head-Control | *trans* | 334 | 320 |  |  | 33.4 (18.3-66.4) |
| Head-Copper | *cis* | 1879 | 1870 | 2152 | 97% | 43.6 (23.9-78.1) |
| Head-Copper | *trans* | 351 | 343 |  |  | 34.2 (20.5-67.4) |
| Head-Response | *cis* | 34 | 34 | 197 | 83% | 34.3 (21.3-64.3) |
| Head-Response | *trans* | 235 | 167 |  |  | 30.0 (18.8-37.5) |

For each of the Control and Copper datasets, *cis* eQTL outnumbered *trans* eQTL (Table 2, Figure S7). In both Response datasets, we identified fewer eQTL overall but *trans* eQTL were more common than *cis* eQTL (Table 2, Figure S7). This pattern is anticipated because the Response datasets are based on the difference in gene expression between control and copper treatments, which should eliminate genetic effects on expression that are consistent in both treatments. Since the bulk of treatment-specific eQTL are *cis* eQTL, the Response data set is expected to have fewer eQTL with local effects.

Across all datasets, the percent variance in gene expression explained by individual eQTL ranged from 10.4% - 78.1% (Figure S10, Table 2). Consistent with previous work (Dixon *et al*. 2007; Emilsson *et al*. 2008; King *et al*. 2014; Albert *et al*. 2018; Keele *et al*. 2020), *cis* eQTL tended to have higher percent variance estimates compared to *trans* eQTL in each of the six datasets (Figure S10, Table 2), suggesting that *cis* eQTL tend to have a larger effect on transcriptional variation compared to *trans* eQTL (Gibson and Weir 2005). However, percent variance estimates are likely overestimated due to Beavis effects so should be assessed with care (Beavis 1995; King and Long 2017).

#### Tissue specificity of mapped eQTL

Although most genes had only one eQTL per dataset (Table 2), 53.7% of genes had one or more eQTL in at least two datasets. For instance, expression of the extracellular Copper/Zinc superoxide dismutase *Sod3* was associated with a single eQTL per dataset but a similarly-located *Sod3 cis* eQTL was detected in five of the six datasets (Figure 4A, B). For other genes, eQTL in different datasets were clearly distinct. For example, expression of the metallothionein gene *MtnA* was associated with a *trans* eQTL in the Head-Copper dataset and a *cis* eQTL in the Gut- Copper dataset (Figure 4C, D). This implies that tissue-specific variants influence *MtnA* expression under copper stress.

**Figure 4.**
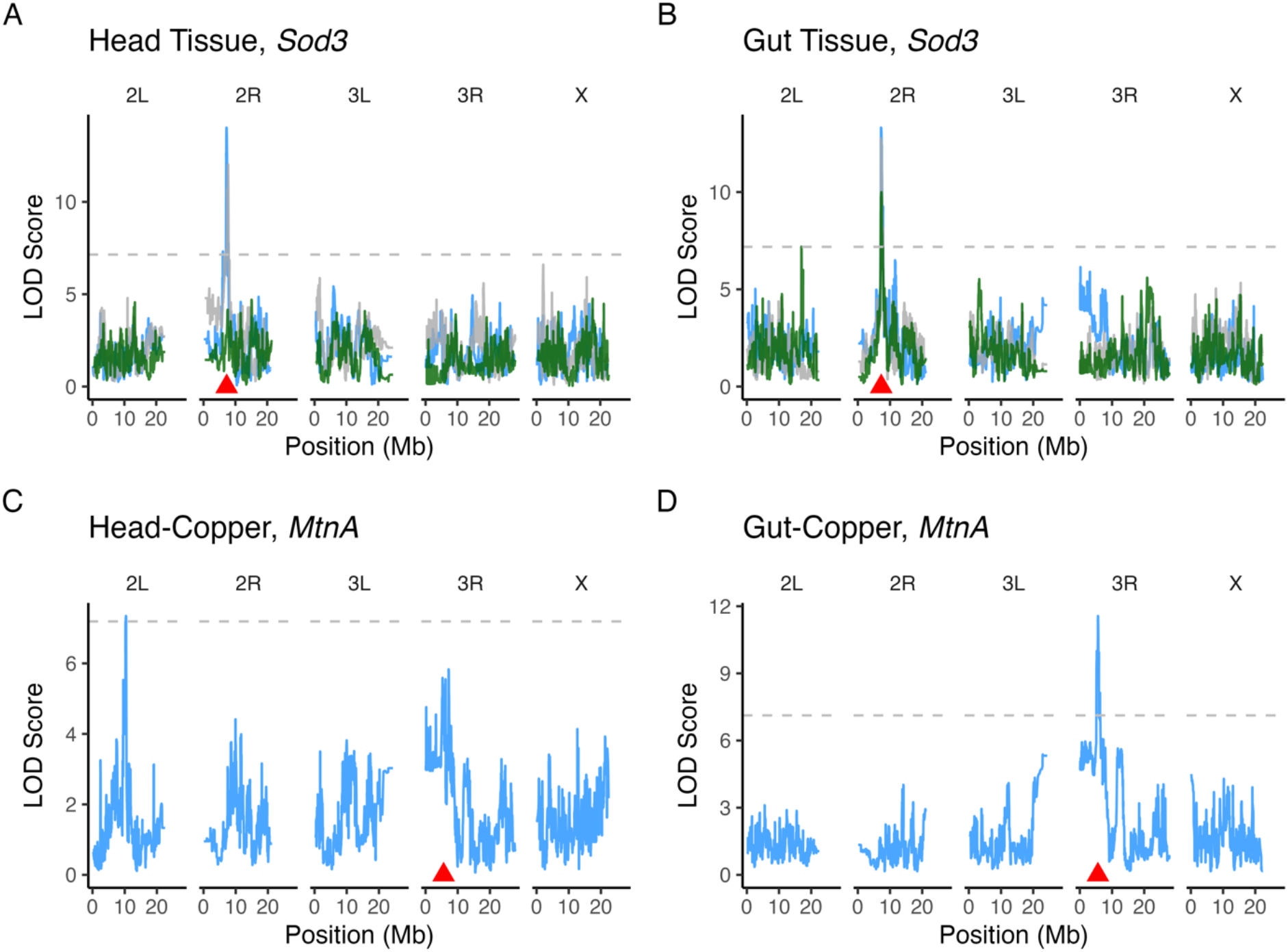
Many genes (53.7%) were associated with eQTL in multiple datasets, but these eQTL were often localized to distinct intervals. A. and B. Expression of the gene Sod3 was associated with cis eQTL in all datasets except Head-Response. C. and D. The gene MtnA was associated with a trans eQTL on chromosome arm 2L in the Head-Copper dataset (C) and was associated with a cis eQTL in the Gut-Control dataset (D). In all plots, the dashed horizontal line indicates the significance threshold based on permutation and the red triangle point indicates the position of the gene. LOD curves are colored based on treatment: gray = Control, blue = Copper, green = Response.

To determine whether a similarly localized eQTL was detected in multiple datasets, we examined eQTL overlap on a per gene basis between tissues within treatment (Head-Control vs Gut-Control, Head-Copper vs. Gut-Copper, Head-Response vs. Gut-Response) and within tissue between treatments (Heads-Control vs Heads-Copper and Gut-Control vs Gut-Copper). eQTL were classified as overlapping if the peak positions were within 1.5cM or if the peak intervals overlapped (see Methods). Overall, *cis* eQTL were more likely than *trans* eQTL to be detected in multiple datasets regardless of which datasets were compared (Figure S11). The number of eQTL that were detected in more than one dataset was highest in within-tissue comparisons (Figure S11A, B), whereas more eQTL were distinct between tissues within treatment (Figure S11C, D). eQTL detected in the Response datasets were most distinct with only 7 *cis* eQTL and no *trans* eQTL shared between Head- and Gut-Response datasets (Figure S11E). Together, these results are consistent with our DE analysis, which identified tissue as having the greatest impact on expression variation (Figure 1).

Within each tissue the majority of *cis* Response eQTL (64-85%) were also detected in the Control and Copper datasets while the majority of *trans* Response eQTL were unique to the Response dataset (Figure S11F-I). The overlapping *cis* Response eQTL were typically detected as *cis* eQTL in both Control and Copper datasets, suggesting that these *cis* eQTL are either associated with different variants in the Control and Copper datasets or that the variant has different additive or magnitude effects on the expression of genes under control and copper conditions. Thorough tests of this hypothesis are beyond the scope of our data; however, correlations between Response and Control or Copper founder haplotype effects at each eQTL suggest both patterns may contribute (Figure S12).

Overall, our eQTL mapping results suggest that regulatory variation influences gene expression in a tissue-specific manner for 2581 genes with distinct eQTL (detected in only one dataset) that were involved in a broad range of processes spanning response to stimulus to cellular metabolism (Figure S13). In contrast, regulatory variants influenced expression of 705 genes with shared eQTL that were detected in both head and gut tissues following copper exposure (Figure S13). These 705 genes were enriched for a smaller number of GO categories including metal response, detoxification, oxidative stress response, and similar stress response pathways (Figure S13). This observation suggests that the control of some major components of metal stress response is potentially consistent across tissues. This said, our data provide some evidence that founder haplotype effects at shared eQTL are not always consistent, suggesting that physically-overlapping eQTL do not necessarily represent identical genetic effects. For example, founder haplotype effects at shared *cis* eQTL in Head and Gut tissue associated with *Mdr49*, *Mdr65*, and *Sid* (involved in insecticide resistance (Sun *et al*. 2017; Denecke *et al*. 2017) and oxidative stress response (Seong *et al*. 2014)) were negatively correlated, implying the regulatory variant may have opposite effects on expression of these genes in head and gut tissue, or that the eQTL actually represent the effects of distinct variants (Figure S14).

We also assessed tissue-specific eQTL patterns using our differential expression analysis results as a guide. Of the 778 stress-related genes (defined using GO terms available from FlyBase) with a tissue by treatment interaction (Figure 3), 349 were associated with a mapped eQTL. Of the 349 genes, 269 had eQTL that were detected in up to three of the four main datasets (for instance, detected in head tissue under copper and control conditions but not in either gut dataset), indicating that most of the genes that were highlighted by DE analysis also had eQTL that followed a tissue- and treatment-specific pattern. Combining our DE and eQTL mapping results provides an additional layer of insight into how gene expression varies across tissue and treatment. For the 269 genes with differential expression and eQTL, allelic variation that influences gene expression variation likely contributes to overall tissue-specific responses to copper stress.

#### Treatment specificity of eQTL

We directly compared the Control and Copper eQTL mapping results within each tissue to identify genetic variants with treatment-dependent regulatory effects on gene expression as well as eQTL with regulatory effects on expression that are consistent over treatments. Overall, we found that eQTL detected under both control and copper conditions were common in head and gut tissue (Figure S11A, B). These overlapping eQTL were associated with expression of genes that included those related to metal response, toxicity response, and copper stress, which resulted in enrichment of gene ontology terms related to stress responses that are consistent with the effects of heavy metal stress (Figure S15). Interestingly, while copper treatment changed the level of expression of metal response genes (based on GO annotation from FlyBase) with eQTL shared by both treatments (Figure 5A, D), the estimated founder haplotype effects at eQTL were generally consistent under both copper and control conditions (Figure 5B, E), i.e., the genetic effects on expression are largely preserved over treatment. Despite this, eQTL had larger effects on gene expression under copper conditions relative to control conditions in both tissues (Head Tissue: F_3,1496_ = 529.2, P < 0.0001, Figure 5C; Gut Tissue: F_3,1258_ = 311.2, P < 0.0001, Figure 5F), and this pattern was most pronounced for eQTL associated with metal response genes in gut tissue (Head Tissue, metal responsive vs non-metal responsive interaction: P = 0.4, Figure 5C; Gut Tissue, metal responsive vs non-metal responsive: P < 0.05, Figure 5F). Therefore, when genes have eQTL in both copper and control treatments, the eQTL contributes to expression variation in each environment in a similar way, but with the magnitude of the eQTL effect varying over tissue.

**Figure 5.**
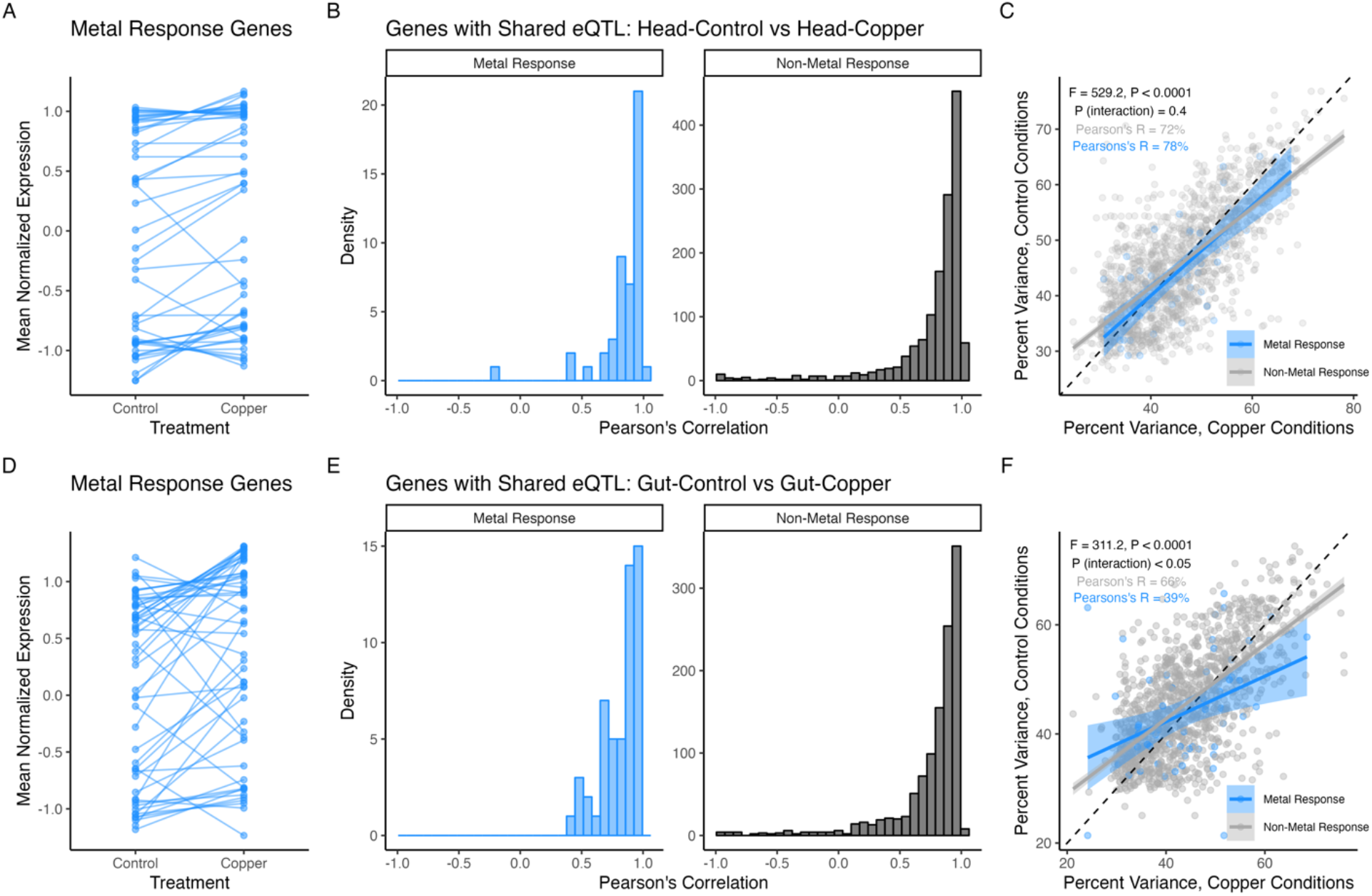
eQTL shared between control and copper conditions are influenced by regulatory variants in a consistent manner across treatments. A. and D. Genes previously linked to copper response, toxicity response, and oxidative stress (metal response genes) shifted in expression level in response to copper treatment in head (A) and gut (D) tissue. B. and E. Founder haplotype estimated effects for shared eQTL were similar under copper and control conditions for both metal response (blue) and non-metal response genes (grey) in head (B) and gut (E) tissues. C. and F. Percent variance explained by shared eQTL estimated in control and copper conditions was positively correlated but indicated larger effects on expression under copper conditions (C, Head Tissue: F_3,1496_ = 529, r^2^ = 51%, P < 0.0001; F, Gut Tissue: F_3,1258_ = 312.2, r^2^ = 43%, P < 0.0001). We found an interaction between percent variance estimates under control and copper conditions for metal response vs non-metal response gene in gut tissue, suggesting that the additive shift in effect size was greater for metal response gene expression variation under copper conditions. In both C and F the dashed line indicates the 1:1 slope to illustrate the reduced slopes of the regressions. Solid lines indicate the best fit using glm estimation and shading indicates the 95% confidence interval of the regression.

eQTL detected in just one of the two treatment conditions per tissue represent potential loci with genotype by environment effects on gene expression. Compared to shared eQTL, there were fewer distinct eQTL between the copper and control treatments for head and gut tissue (Figure S11A, B). In head tissue, we found copper-specific eQTL associated with genes enriched for functions related to ion, cation, and metal ion transport (Figure S16A). Copper-specific head eQTL were associated with several copper-related genes including *MtnA*, *Mco1*, *Mco4*, *Loxl2*, and *CG17996* (Thurmond *et al*. 2019; The UniProt Consortium *et al*. 2021). Of note are *MtnA* and *Mco1*. *MtnA* encodes one of five metal scavenger proteins that reduce the levels of free copper ions available in the cell (Calap-Quintana *et al*. 2017). Exposure to copper increases *MtnA* expression in head tissue (Figure 2B), and the *trans* eQTL associated with *MtnA* (Figure 4C) may indicate a distant regulatory element that influences expression under copper stress. *Mco1* uses copper as a cofactor for iron metabolism (Lang *et al*. 2012) and was associated with a *cis* and a *trans* eQTL. The *trans* eQTL associated with *Mco1* falls near the metal-dependent transcription binding factor *MTF-1*, which is one of the primary transcription factors involved in metal stress response (Egli *et al*. 2003; Balamurugan *et al*. 2004). Control-specific eQTL in head tissue highlighted a set of genes primarily enriched for functions related to broad categories including cellular process and response to stimulus (Figure S16B). Despite the lack of enrichment for metal-related GO terms, control-specific eQTL were associated with several genes involved in copper ion homeostasis including *Mvl*, *CG10505*, *CG7194*, Grx1, and *Ctr1B* (Thurmond *et al*. 2019; The UniProt Consortium *et al*. 2021). Maintaining proper copper homeostasis is critical for normal physiological function, and these eQTL with effects on expression of copper homeostasis genes may provide insight into regulation of copper transport under non-stressful conditions.

In gut tissue, we found enrichment for GO categories related to lipid metabolism, localization, and homeostasis among genes with copper-specific eQTL (Figure S17A) and enrichment for GO categories related to localization, transport, and regulation among genes with control-specific eQTL (Figure S17B). Two copper-related genes (*olf413* and *CG6908*) had eQTL under copper conditions in gut tissue (Thurmond *et al*. 2019; The UniProt Consortium *et al*. 2021). Both eQTL were far from the gene positions (*trans*) and neither were near known metal-responsive transcription factors. Copper-specific eQTL also implicated several genes from the glutathione s transferase and cytochrome p450 families of genes, which are involved in the response to toxic substances (Kim and Yim 2013; Esteves *et al*. 2021), and we observed a copper- specific eQTL for the metallothionein gene *MtnA*. Similar to head tissue, several copper homeostasis genes had eQTL that were only detected under control conditions including *Ccs*, *CG11590*, *CG11825*, and *Atox1* (Culotta *et al*. 1997; Hatori and Lutsenko 2013), suggesting that allelic variation in the DSPR contributes to variation in regulation of expression of these genes under non-stressful conditions.

Treatment-dependent eQTL were also identified using the Head- and Gut-Response datasets. Response eQTL reveal loci that impact the difference in gene expression between copper and control conditions in a genotype-specific manner and are thus inherently genotype by environment eQTL. Response eQTL detected in both Gut- and Head-Response datasets were associated with several potential candidate genes that may play a role in the response to copper toxicity. GO analysis of genes with Gut-Response eQTL highlighted categories related to detoxification pathways and response to toxins including glutathione s transferase family genes (glutathione metabolic process; hypergeometric test for overrepresentation of GO terms, adj p < 0.001; Figure 6) and cytochrome p450 genes (response to toxic substances and insecticides, adj p < 0.001; Figure 6) (Kim and Yim 2013; Esteves *et al*. 2021). We also detected Response eQTL for two ABC transporter multidrug resistance protein genes (*Mdr49* and *Mdr69*) as well as two genes that are known to play a role in copper metabolism or binding (*Sod3* and *Tbh*) (Itoh *et al*. 2009; The UniProt Consortium *et al*. 2021). More generally, 50 out of 316 annotated genes with Response eQTL in the Gut-Response dataset are known or predicted to be involved in metal ion binding (Thurmond *et al*. 2019; The UniProt Consortium *et al*. 2021).

**Figure 6.**
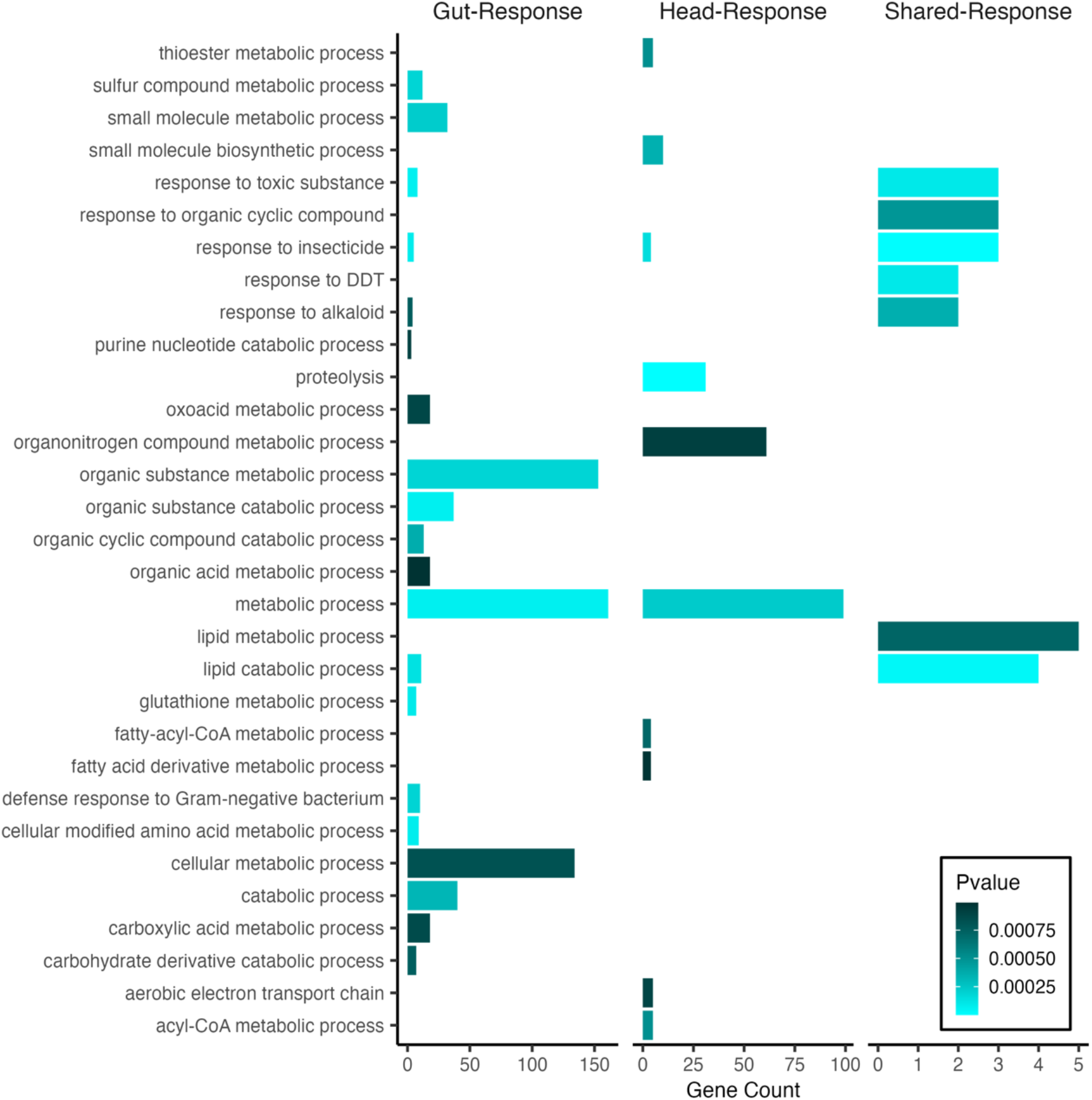
Enrichment of GO categories for Response eQTL genes by tissue (head tissue = 197 genes; gut tissue = 316) and for the set of eQTL genes that were shared between tissues (Shared-Response, 19 genes).

Head-Response eQTL included fewer potential candidate genes. Top GO enrichment categories included proteolysis and metabolic process (adj p < 0.0001) followed by response to insecticide (adj p < 0.001; Figure 6). We detected Head-Response eQTL for ABC transporter gene *Mdr50* as well as a different *trans* eQTL associated with *Mco1* that was distinct from those identified by comparing the copper and control-specific head eQTL. Of the 197 annotated Response eQTL genes, 26 play or are predicted to play a role in metal ion binding. There were 19 eQTL genes that were shared between the Head- and Gut-Response datasets, which included several cytochrome p450 family genes. The shared set of Response eQTL were associated with expression of genes that contribute to response to insecticide (adj p < 0.0001), response to toxic substance (adj p < 0.001) and response to DDT (adj p < 0.001) (Figure 6).

#### Identification of eQTL hotspots

Using a chi-square goodness of fit test, we identified between two and seven chromosomal regions that encompassed more *trans* eQTL peaks than expected by chance per dataset; these are potential *trans* eQTL hotspots (Table 3). Head- and Gut-Response datasets had the most hotspots, and overall, more hotspots were detected in head tissue compared to gut tissue datasets (Table 3). Several hotspots flanked the centromeric regions of chromosomes 2 and 3 (Figure 7) and were detected in multiple tissues and datasets (Table 3; Figure 7). This is likely because we identified hotspot locations using genetic distance, and since crossing over is low and gene density is higher in centromeric regions, the chance of detecting a hotspot at centromeres is likely inflated.

**Figure 7.**
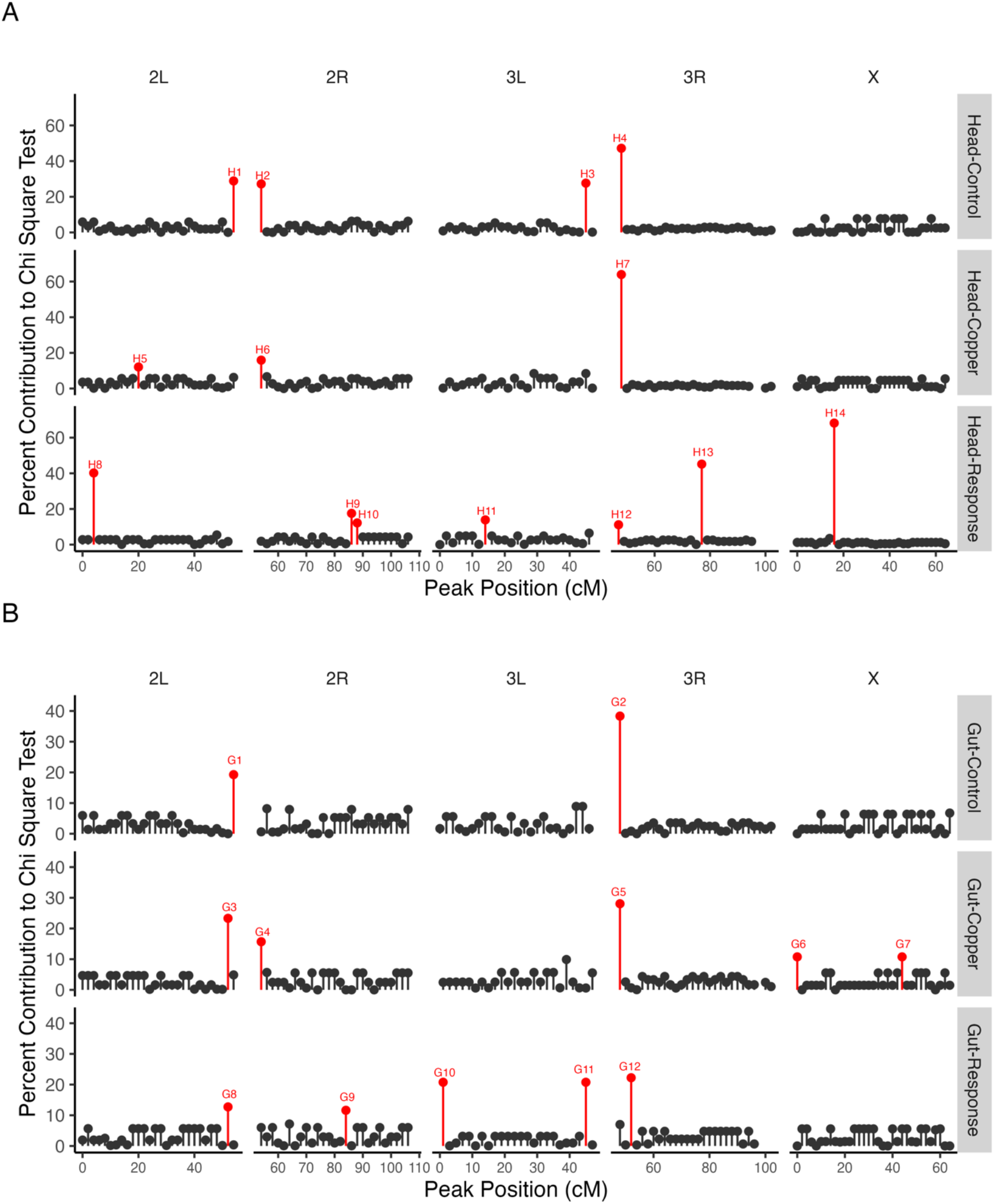
We detected enrichment for trans eQTL in all datasets and tissues. Several hotspots (primarily near centromere regions) were detected in multiple tissues and datasets. The majority of trans eQTL hotspots were detected in Response datasets in both head and gut tissue.

**Table 3.**
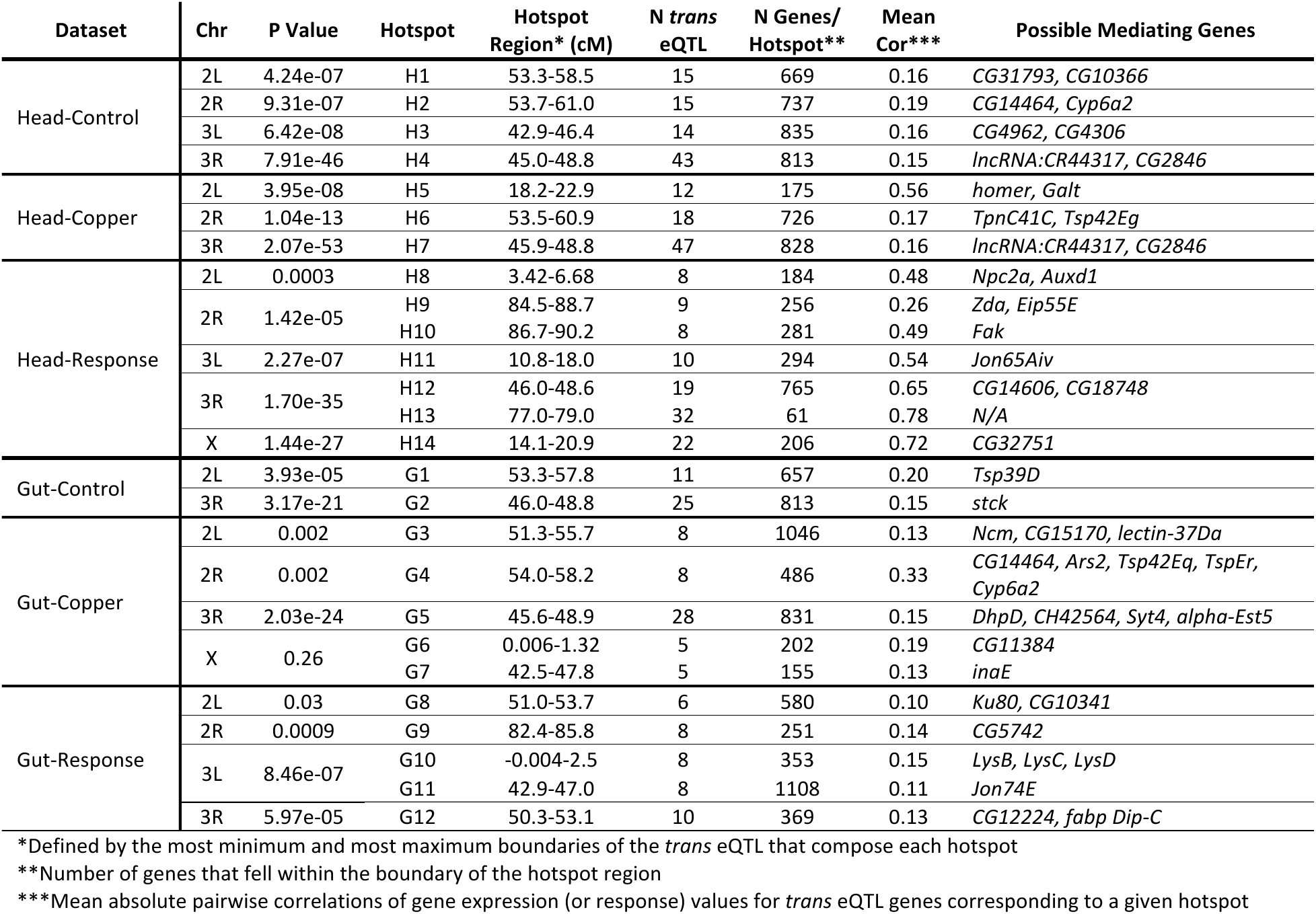
Chi-square analysis of trans eQTL distribution along chromosome arms and characteristics of hotspots.

If all genes with a *trans* eQTL in a given hotspot are co-regulated by a genetic element in the hotspot region, we anticipate that the expression of these genes (or their expression response to copper treatment) will be correlated to some extent. Mean absolute pairwise correlations of gene expression or gene response values for *trans* eQTL genes corresponding to a given hotspot ranged from 6-78% in Head datasets and from 10-60% in Gut datasets (Figure S18, Figure S19; Table 3). We performed PCA on the matrix of per-RIL expression or response measures for all the genes with a *trans* eQTL in each hotspot, with PC1 representing a composite variable of the shared expression (or response) variation of genes with *trans* eQTL in that hotspot. All composite hotspot variables mapped a QTL at the location of the hotspot, and genes with high pairwise correlations (Figure S18, Figure S19) had consistent loadings on PC1 in each hotspot (Figure S20).

To identify genes that may modulate expression of the set of genes with *trans* eQTL at a hotspot, we used a mediation analysis approach that treated the expression level of each gene that was encompassed by a hotspot interval (Table 3) as a covariate in QTL analysis of the composite hotspot variable (see Methods). Including a true mediator as a covariate in the analysis is expected to eliminate the presence of the composite variable QTL, resulting in a drop in the LOD score at the peak position. With this approach we were able to identify between one and five genes that had complete or partial mediating effects on the composite variable for all but one hotspot (Figure S21, Figure S22; Table 3).

Our mediation analysis revealed three candidate genes that completely accounted for the H11 composite hotspot QTL (Figure 8A). H11 is composed of 10 *trans* eQTL peaks that were detected in the Head-Response dataset (Table 2). The three candidate mediating genes (*Jon65Aiv*, *Jon65Aiii*, and *yip7*) are serine proteases expressed in the adult head and are involved in proteolytic activity and protein metabolism (Kumar *et al*. 2021) (Figure 8A). We did not identify any eQTL associated with *Jon65Aiii* or *yip7* but did detect one Head-Response *cis* eQTL near *Jon65Aiv* and compared the estimated haplotype effects at this QTL with those at the H11 composite peak. Estimated founder haplotype effects were significantly negatively correlated (r = -99%, p < 0.001, Figure 8B), suggesting that the haplotypes at this location have similar effects on the response of *Jon65Aiv* to copper and on the response at genes that have a response *trans* eQTL at the H11 hotspot. *Jon65Aiv* was recently identified as a novel candidate gene that may contribute to the gene expression response to copper stress in *D. melanogaster* strains derived from wild populations (Green *et al*. 2022). Copper exposure in wild-collected flies from that study resulted in decreased expression of *Jon65Aiv*; we did not observe a significant change in expression of any of the three H11 candidate mediating genes in response to copper exposure.

**Figure 8.**
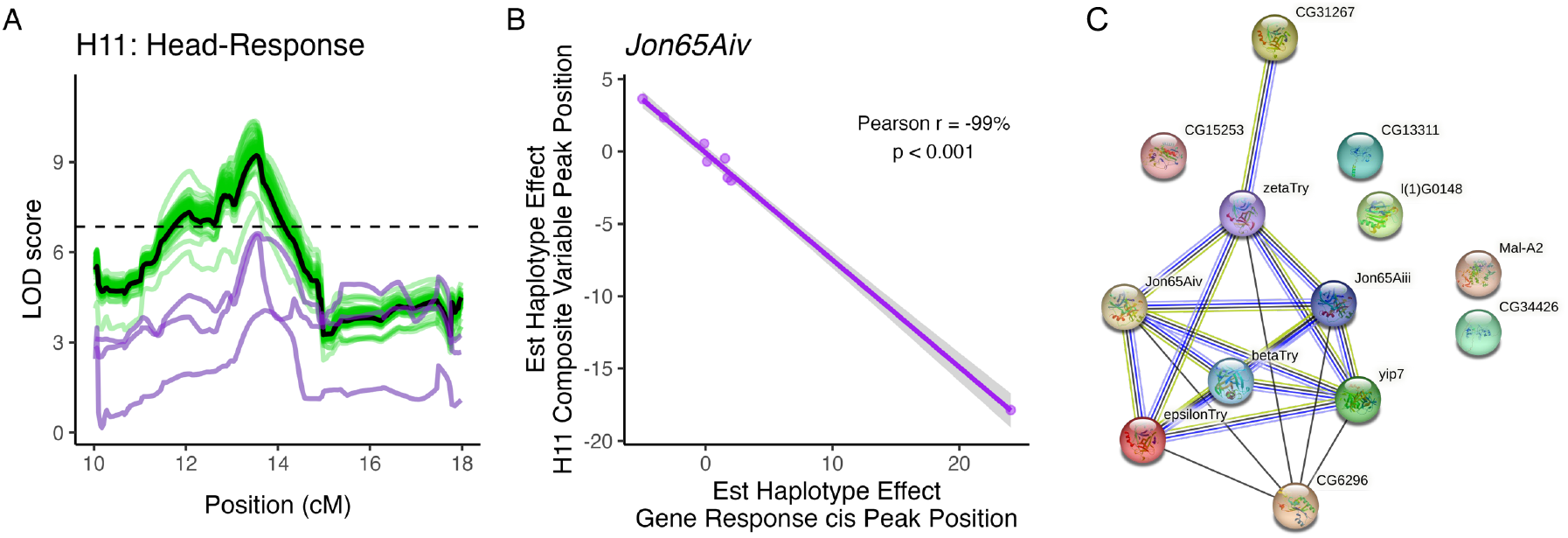
Three genes had complete mediating effects on the composite eQTL peak for hotspot H11. A. Including Jon65Aiv (bottom purple line), Jon65Aiii, and yip7 (top purple lines) individually as covariates accounted for the composite QTL peak for H11 (black line). No other gene encompassed by the H11 composite peak had a strong mediating effect on the composite peak (green lines). B. Estimated founder haplotype effects at the composite QTL peak and at a cis eQTL associated with Jon65Aiv were significantly negatively correlated (Pearson r = -99%; p < 0.001). C. STRING analysis of the three candidate mediating genes and genes with a trans eQTL peak at the hotspot suggests that the candidate mediating genes are co-expressed with trans eQTL genes rather than having a regulatory role. Black edges indicate co-expression, green edges indicate similarity through textmining, light blue edges indicate protein homology, and dark blue edges indicate gene co-occurrence.

We leveraged the STRING database to examine potential interactions between candidate mediating genes and genes with *trans* eQTL at the H11 hotspot (Szklarczyk *et al*. 2023). Using information gathered on predicted and known protein interactions and gene co-expression, STRING analysis showed that *Jon65Aiv*, *Jon65Aiii*, and *yip7* are directly or indirectly co-expressed with five of the genes with *trans* eQTL at the H11 hotspot, and several of the H11 *trans* eQTL genes have serine-type endopeptidase activity (GO enrichment analysis; p < 0.001), protein catalytic activity (p < 0.001), or hydrolase activity (p < 0.001) (Figure 8C). Together, these results suggest that *Jon65Aiv*, *Jon65Aiii*, and *yip7* are not regulatory mediating genes but rather in a shared pathway with the other H11 *trans* eQTL genes.

We also identified a serine protease as a candidate gene with a partial mediating effect on the Gut-Response G11 *trans* eQTL hotspot (Figure 9A). *Jon74E* response to copper was associated with a *cis* eQTL, and similar to *Jon65Aiv*, estimated founder haplotype effects of the *cis* eQTL were negatively correlated with the estimated haplotype effects at the composite G11 hotspot peak (Pearson’s r = -98%, p < 0.001, Figure 9B). Unlike the previous candidate mediating genes, *Jon74E* did not have any connections to any other gene with a *trans* eQTL at the G11 hotspot (Figure 9C).

**Figure 9.**
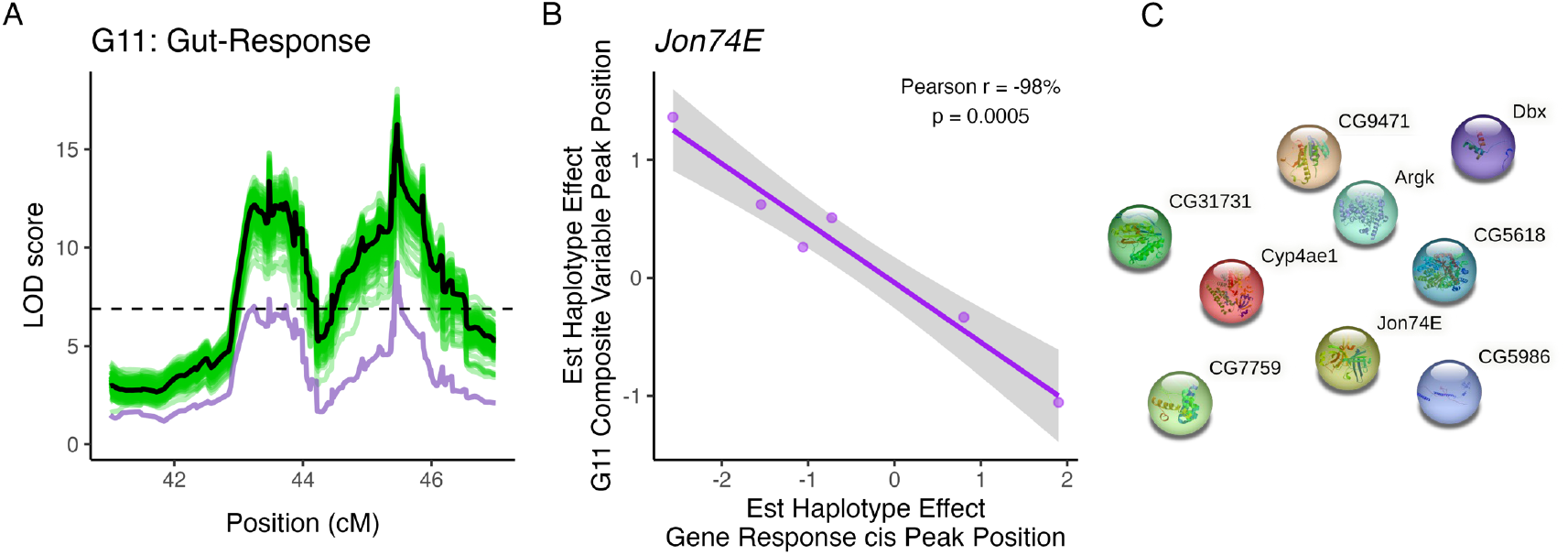
One gene had a partial mediating effect on the composite eQTL peak for hotspot G11. A. Including Jon74E (purple line) as a covariate in QTL analysis of the G11 composite variable accounted for the composite QTL peak for H11 (black line). No other gene encompassed by the G11 composite peak had a strong mediating effect on the composite peak (green lines). B. Estimated founder haplotype effects at the composite QTL peak and at a cis eQTL associated with Jon74E were significantly negatively correlated (Pearson r = -98%; p < 0.001). C. STRING analysis of the candidate mediating gene and genes with a trans eQTL peak at the hotspot did not provide any insight into the link between the candidate mediator and the genes with trans eQTL in the hotspot.

Overall, while we identified several genes with expression patterns that appeared to mediate a *trans* eQTL hotspot effect (Figures S21 and S22), we found limited evidence of functional links between expression of candidate mediating genes and the genes with *trans* eQTL at the corresponding hotspot. Our hotspot analysis did not identify any candidate transcription factors or genes that have previously reported regulatory effects on *trans* eQTL genes in either head or gut tissue.

## Discussion

### Copper exposure results in tissue- and treatment-specific expression

The genetic architectures of complex traits such as stress resistance and disease are highly context-dependent and can vary across scales as broad as between species and populations (Whitehead and Crawford 2005; Breschi *et al*. 2016; Findley *et al*. 2021) and as small as between individuals, regions of a particular tissue (Fournier and Schacherer 2017; Çelik and Akdaş 2019; Munro *et al*. 2022), or even within individuals across age or development (Everman and Morgan 2017; Huang *et al*. 2020; Everman *et al*. 2021). Given anticipated tissue-specific gene expression patterns resulting from the spatial distribution of specialized copper-accumulating cells (Calap-Quintana *et al*. 2017; Miguel-Aliaga *et al*. 2018) and the potential for copper toxicity to result in acute damage to digestive tissues as well as neurological tissues (e.g., Tchounwou *et al*. 2008; Jomova *et al*. 2010), one of our principal goals was to characterize the tissue-specific transcriptomic response to copper toxicity. We demonstrate striking patterns of tissue- and treatment-specific genetic response to copper exposure using a combination of differential expression analysis and eQTL mapping. More than 90% of all genes had tissue-specific patterns of expression, and a significant tissue by treatment interaction affected 70% of the genome (Figure 1). Our data suggest that this interaction may be due to greater magnitude and diversity of the gene expression response in Gut tissue compared to Head tissue (Figure 3). Particularly, when stress-related genes were considered as a group we found that the stress response in gut tissue involved multiple stress pathways, whereas oxidative stress response was the primary stress response pathway activated in head tissue (Figure 3). Our expression QTL (eQTL) mapping results provide similar insight; the majority of eQTL were tissue-specific within treatment (Figure S11), suggesting that the genetic control of gene expression response to stress is highly context-dependent.

In general, overexposure to heavy metals results in the accumulation of reactive oxygen species (ROS) and activation of oxidative stress response pathways (e.g. Ercal *et al*. 2001; Gaetke 2003; Uriu-Adams and Keen 2005). Especially for heavy metals that are biologically necessary, there are also metal-specific metabolism pathways that can contribute to the metal toxicity response (Calap-Quintana *et al*. 2017). In the case of copper, specialized cells that are involved in uptake, metabolism, and detoxification of copper ions from the diet line the middle midgut of the fly and the acidic region of the digestive system in vertebrates (Dubreuil 2004; Calap-Quintana *et al*. 2017). Many of the genes that have been linked to copper response in previous studies were significantly differentially expressed under copper conditions relative to control conditions in our study (Figure 2), and our eQTL mapping results suggest that many of these copper-responsive genes are influenced by genetic variation in regulatory elements. Furthermore, we found that genes with functions related to metal response followed expected tissue-specific expression patterns (e.g. *Mtn* family genes; *Ctr1A* and *Ctr1B*; *Atox1* and *SOD3*; *ATP7*; discussed above (Zhou *et al*. 2003; Turski and Thiele 2007; Itoh *et al*. 2009; Calap-Quintana *et al*. 2017; Kamiya *et al*. 2018)) (Figure 2). For instance, and consistent with previous reports, we found that *MtnA* expression was strongly induced by copper exposure (Figure 2B); however, the level of induction was much more pronounced in gut tissue compared with head tissue, leading to a significant tissue by treatment interaction. We also found variation in the genetic control of *MtnA* expression under copper conditions that followed a tissue specific pattern. In gut tissue, *MtnA* expression in response to copper is influenced by a *cis* eQTL (Figure 4C), whereas in head tissue *MtnA* expression under copper conditions is influenced by a *trans* eQTL (Figure 4D). Although the genetic variants that contribute to *MtnA* expression in either tissue are present in both tissues, each variant appears to only influence gene expression in a particular tissue.

While *D. melanogaster* is commonly used to characterize the genetic control of heavy metal response (e.g. Egli *et al*. 2003, 2006a; Balamurugan *et al*. 2004; Norgate *et al*. 2010; Hua *et al*. 2011), tissue-specific comparisons are not common (Fasae and Abolaji 2022). One of the few examples of tissue-specific response to heavy metal toxicity in *D. melanogaster* demonstrated that variation in *CncC* pathway activity (involved in autophagy regulation) across muscle and neurological tissue contributed to the toxicity response to methylmercury (Gunderson *et al*. 2020). Our differential expression analysis provides novel insight and suggests that there are tissue-specific pattens in broad responses to stress through the differential activation of pathways. Although all the stress response categories represented by our subset of genes can be linked to heavy metal stress response (Darling and Cook 1843; Kefaloyianni *et al*. 2005), our data suggest that oxidative stress response is the primary stress response mechanism to copper exposure in neurological tissue (proxied with head tissue in our study) while the primary stress response mechanisms in gut tissue are more diverse. Because previous studies that offer insight into tissue-specific gene expression response often focus on sets of tissues that infrequently overlap, it is challenging to determine whether the patterns we observe in our study are unique or reflective of a general pattern. However, our data do strongly suggest that there is a tissue- specific response to copper toxicity over the eight-hour exposure period used in this study. Whether this difference is due to tissue-specific vulnerability, to mode of exposure, to temporal progression of copper ions throughout the organism, or a combination of these and other factors remains to be fully determined. Additional follow up with tissue-specific knockdown or ablation of key genes involved in oxidative stress and other stress response pathways would be needed to strengthen these observations.

### Transcriptomic response to copper stress is influenced by regulatory allelic variation

Allelic variation and its contribution to trait variation has been repeatedly dissected, characterized, and functionally tested in the context of complex traits ranging from those associated with human disease (e.g., Granhall *et al*. 2006; Hardy *et al*. 2018) to various stress responses including heavy metal resistance (e.g., Zhou *et al*. 2017; Evans *et al*. 2018; Everman *et al*. 2019, 2021). In addition to the potential to influence the functional gene product, allelic variation can also influence the response to stress via effects on gene expression patterns. The majority of allelic variation is found in non-coding regions of the genome (Umans *et al*. 2021), leading to the expectation that these non-coding variants play a regulatory role. By treating expression of individual genes as traits measured in multiple treatments and tissues, we were able to gain insight into context-dependent genetic control of gene expression using expression eQTL mapping. Approximately half of the annotated *Drosophila* genome was associated with one or more eQTL in our study, providing ample evidence of genetic variation in the control of gene expression (Table 2). Nearly all (98%) of genes that were associated with at least one eQTL were also differentially expressed, suggesting that the variants associated with these genes likely influence expression, although functional validation would be necessary to verify this hypothesis. This observation is consistent with work by Boyle *et al*. (2017) demonstrating that SNPs which contribute to trait variation are enriched near genes that are actively transcribed under disease states; our observations suggest this pattern extends to stress conditions as well.

Although eQTL mapping provides detailed insight into the genetic control of a given trait by examining elements that contribute to trait variation on a gene-by-gene basis, there are several challenges with this approach. Power to detect eQTL with modest effects is directly related to the number of strains in which the trait is measured (Ruden *et al*. 2009; King *et al*. 2012a; Qu *et al*. 2018; Arvanitis *et al*. 2022). By using 93-96 genotypes in our eQTL analyses, lack of power to detect modest-effect eQTL likely contributes to our estimates of tissue-specificity. For example, power estimates to detect a QTL with an effect size of 10 with 100 DSPR strains is approximately 20% (King *et al*. 2012a). Detection of tissue-specific eQTL is also more sensitive to false negatives, for example due to low levels of expression of a particular gene in a given tissue. However, despite these challenges our data suggest that allelic variation that contributes to the regulation of gene expression is tissue-specific for approximately 65% of eQTL detected in the control and copper datasets.

Similar studies of tissue-specific eQTL mapping are relatively rare in model organisms but have provided examples of tissue-specific eQTL. Using a mouse multiparental mapping population (Collaborative Cross (Aylor *et al*. 2011)) that is similar in concept to the DSPR, Keele *et al*. (2020) demonstrated that among lung, liver, and kidney tissue genetic control of several genes varied across tissues. Spatially discrete genetic control of gene expression can also vary at the sub-tissue level. Munro *et al*. (2022) demonstrated gene expression patterns that are specific to five regions of the rat brain are influenced by a combination of shared and sub-tissue specific eQTL. Shared eQTL were most common in similar brain tissue types, but sub-tissue specific eQTL were common between more distinct types of brain tissue (Munro *et al*. 2022). eQTL mapping studies of human disease and native state gene expression have also provided evidence of tissue- and cell type-specific genetic control of gene expression (Dimas *et al*. 2009; Nica *et al*. 2011; Fairfax *et al*. 2012; Peters *et al*. 2016; GTEx Consortium 2017; GTEx Consortium *et al*. 2018). A recent study presenting a novel analytical approach to identify colocalized eQTL suggests that tissue-specific eQTL may be more common than previously thought and key for identifying regulatory variants associated with complex disease traits (Arvanitis *et al*. 2022). The existence of other examples of tissue-specific eQTL combined with our evidence strongly suggest that tissue-specific eQTL contribute to the stress response to copper toxicity in *D. melanogaster*.

### Consistent versus tissue-specific effects of eQTL

In addition to identifying tissue-specific eQTL, our study also provides insight into eQTL that have consistent versus environment-dependent effects within tissue type. Complex traits are dependent on both genotype and environmental variation, and the interaction between genotype and environment can contribute to overall susceptibility of individuals to stress and disease (Li *et al*. 2006; Duveau and Félix 2012; Moyerbrailean *et al*. 2016; Lea *et al*. 2022). The importance of genotype by environment interactions and the detection of quantitative loci with context-dependent effects was demonstrated by Lea *et al*. (2022) in a study using 544 human cell lines exposed to 12 environments. Using a high-powered experimental design, they demonstrated that the complex traits measured were influenced by a combination of eQTL with consistent effects in multiple environments as well as eQTL with environment-dependent effects (Lea *et al*. 2022). Similarly, Umans *et al*. (2021) offer a review and perspective on the importance of examining regulatory variants in multiple contexts. Recognizing the dynamic nature of context-dependent effects of alleles associated with disease in humans, Umans *et al*. (2021) propose that a critical approach to characterizing allelic variation in regulatory elements is to use multiple treatment conditions to detect important disease associated variants.

Overall, the majority of the eQTL we detected were near the position of the gene they were associated with (*cis* eQTL), suggesting that genetic variation in local regulatory elements plays an important role in the gene expression response to copper stress. *cis* eQTL generally have larger effects on trait variation and are thus easier to detect with modestly powered designs (Hill *et al*. 2021), and *trans* eQTL are additionally difficult to detect because they may not act at the level of mRNA regulation but instead have larger detectable effects at the protein level (Boyle *et al*. 2017). *cis* regulatory variation is widely appreciated to be an important contributor to phenotypic variation and plays a decisive role in the evolution of traits in natural populations and in human disease (Wray 2007; Savinkova *et al*. 2009; Wittkopp and Kalay 2011; Hill *et al*. 2021). At the treatment level and consistent with previous studies in humans, flies, and worms (Ruden *et al*. 2009; Snoek *et al*. 2017; Qu *et al*. 2018; Sterken *et al*. 2020; Lea *et al*. 2022), we found eQTL consistently in both treatments as well as eQTL that were only detected in one of the two treatments (Figure S11). The number of eQTL that were consistently detected was higher than treatment-specific eQTL, similar to a pattern previously reported for human cell lines (Lea *et al*. 2022). Similarly, in *D. melanogaster*, Qu *et al*. (2018) and Ruden *et al*. (2009) demonstrated that a small but non-negligeable number of eQTL were only detected following exposure to lead in head tissue or whole animals, respectively. Our analysis provides an additional layer of insight by examining the correlation between the estimated haplotype effects at each of the peaks that were consistently detected under control and copper treatment conditions in head and gut tissue (Figure 5). We found that estimated effects of founder alleles at shared eQTL often had similar effects on expression of the gene under different conditions. Despite this apparent consistency, an effect of treatment was still present in the eQTL detected under copper and control conditions in gut tissue; eQTL associated with metal-associated genes had larger effects on gene expression variation under copper conditions compared to control conditions (Figure 5F). While our results highlight the potential for even eQTL that are repeatedly detected across treatments to have treatment-specific effects on the expression of a given gene, adding a deeper dimension to gene by environment interaction eQTL, it is important to consider that the QTL intervals may include more than one variant that influences gene expression. One of these variants may influence expression under control conditions while the other influenced expression under copper conditions. Without additional follow-up studies, our analyses do not provide sufficient resolution to distinguish this case from one in which the same variant has treatment-specific effects on gene expression.

In addition to characterizing genotype by environment eQTL by examining the difference between the control and copper datasets for each tissue, we also examined eQTL that were associated with a summary metric of the gene expression response to copper stress. The majority of eQTL detected in the Head- and Gut-Response datasets were *trans* eQTL. Our study design partially accounts for the deficit in *cis* eQTL in the Response datasets, as any effects of *cis* eQTL that were detected with similar effects in control and copper datasets would be reduced in the Response eQTL mapping analyses. However, an enrichment of *trans* eQTL associated with specific environments has been previously observed in *C. elegans* in response to temperature stress (Li *et al*. 2006; Snoek *et al*. 2017). *trans* eQTL that are associated with the dynamic shift in gene expression response to the environment call attention to allelic variants that may play a role in the regulation of stress-dependent expression change that may otherwise not be detected if the treatment conditions are considered independently. We found several Response *trans* eQTL near genes with a wide range of functions related to response to toxic conditions (Figure 6) that highlight potential candidate regulatory QTL. Additional follow up would be necessary to corroborate our observations and to fully characterize the role that these potential QTL sites may play in the regulation of the response to copper stress.

## Conclusions

Exposure to heavy metal stress results in a profound change in transcriptional activity across multiple tissues, and the genetic control of the gene expression response varies depending on the environmental conditions and tissue. By taking a combined approach of differential expression analysis and eQTL mapping, we were able to dissect and characterize more subtle differences in tissue-specific response to copper toxicity. Our work provides a novel set of candidate loci that may have context-dependent effects on gene expression and plasticity. From the patterns that differentiate the genetic response observed in head and gut tissue, we gain deeper insight into the level of toxicity response that is activated by short exposure to copper stress.

## Acknowledgements

We thank Kristen Cloud-Richardson for assistance with collection of head tissue. We also thank the anonymous reviewers for their suggestions that improved this manuscript.

## Funding

This project was supported by funding from NIH: ERE was supported by two postdoctoral fellowships (F32 GM133111 and K99 ES033257) and by the Kansas INBRE (Kansas Idea Network of Biomedical Research Excellence) project (P30 GM145499). Additional support for experiments was provided by NIH R01 ES029922 and R01 OD034064 awarded to SJM. Sequencing was performed by the KU Genome Sequencing Core which is supported by the Center for Molecular Analysis of Disease Pathways and NIH Centers of Biomedical Research Excellence (P30 GM145499).

## Conflict of Interest

The authors declare no conflicts of interest.

## Supplemental Figures

**Figure S1.** Adult copper resistance of B panel RILs used in the current study. We randomly sampled 48 resistant and 48 sensitive DSPR B panel RILs from the top and bottom 25% of the distribution of adult copper survival response following 48-hour exposure to 50mM CuSO_4_ reported in (Everman et al. 2021).

**Figure S2.** Distributions of raw read pair counts by library pool. Each tissue-specific pool included copper and control sample pairs, and an even representation of resistant and sensitive RILs. HO stands for “high-output” NextSeq500 flowcell, MO stands for “mid-output” flowcell. Raw reads from the two sequencing runs of the 96-plex Heads library (green shading) were combined prior to read processing.

**Figure S3.** Comparisons of the number of raw read pairs, the number retained following trimming, the percentage aligned, and the percentage assigned to genes following the HISAT2 pipeline from the high-output (HO) and mid-output (MO) sequencing runs for the 96-plex Head library. Apart from raw pair count, which varies due to under-clustering of the 96-plex Head library, data are consistent between the HO and MO sequencing of the same 96-plex library and are comparable to the separate 94-plex Head library that was only sequenced on one HO flowcell.

**Figure S4.** Variation in read length across all 379 samples. After trimming low quality bases and filtering short reads, only read pairs with >15 bp per read were retained.

**Figure S5.** Gene expression estimates from the HISAT2 and kallisto pipelines were consistent. After alignment and filtering out of lowly-expressed genes, expression estimates for 9842 and 9913 genes from the HISAT2 and kallisto pipelines, respectively, were retained. Gene-wise correlations in read counts across all samples for shared retained genes (9434) were highly correlated (mean R = 98.2%).

**Figure S6.** Correcting for known and unknown sources of technical variation reduced noise in eQTL mapping. Head-Control (top row) and Head-Copper (middle row) eQTL maps for *Sod1* are noisy when technical variation is not accounted for (left column). Correcting for 2% (middle column) and 1% PCs (right column) along with those associated with known technical factors appears to limit noise. Head-Response data (bottom row) were less sensitive to data correction. In each plot, orange vertical lines indicate the location of the gene. Horizontal lines indicate the gene- and dataset-specific significance threshold.

**Figure S7.** Correcting for PCs that explained 2% or 1% of the variation in gene expression resulted in similar numbers of eQTL. Head samples are shown in A; Gut samples are shown in B. In both panels, the left column shows the 2% correction, the right column shows the 1% correction.

**Figure S8.** Genome-wide LOD scores were similar between the 1% and 2% corrected datasets for each tissue and treatment. Head-Copper, -Control, and -Response correlations are shown in A-C. Gut-Copper, -Control, and -Response correlations are shown in D-F. In each panel, orange highlights genome-wide LOD score correlations for genes that are associated with at least one eQTL in the 2% corrected data; grey highlights genes that were not associated with eQTL. Vertical color-coded lines indicate mean correlations for each group of genes.

**Figure S9.** For most genes (>80%), mapping yielded a single eQTL. Data shown are counts of distinct eQTL peaks per gene for each of the 6 datasets. Gut tissue is shown in orange, head tissue is shown in red. Numbers above bars provide the actual number of genes in each category.

**Figure S10.** Percent variance explained by mapped *cis* and *trans* eQTL. *cis* eQTL (blue) tended to have higher estimates compared to *trans* eQTL (red). Estimates were slightly lower in both response datasets for both *cis* and *trans* eQTL. Numbers at the base of each plot report the total number of *cis* and *trans* eQTL detected per dataset.

**Figure S11.** Many eQTL were detected in multiple datasets. Overlap was highest within tissue between treatments (A and B) but was still substantial in comparisons between tissues within treatment (C, D). Comparisons between the Head- and Gut-Response datasets revealed that most eQTL were distinct between the tissue-specific characterization of copper response. Overlap was also high in comparisons of the Response datasets against the tissue- and treatment-specific datasets (F – I). The majority (64 – 81%) of *cis* Response eQTL were shared with either the Control or Copper datasets for the corresponding tissue. For Head tissue, all but 9 *cis* Response eQTL were shared amongst Head-Control or Head-Copper *cis* eQTL; for gut tissue, all but 14 *cis* Response eQTL were shared amongst Gut-Control or Gut-Copper cis eQTL (data not shown).

**Figure S12.** Founder haplotype effects at *cis* eQTL peak positions that were detected in Response, Copper, and Control datasets for a given tissue ranged in strength of correlation for each comparison: A. Head-Response vs Head-Control, B. Head-Response vs Head-Copper, C. Gut- Response vs Gut-Control, D. Gut-Response vs Gut-Copper. Founder effect estimates are impacted by low replication per DSPR strain and should be interpreted with care; however, patterns suggest the detection of Response eQTL in our study may be influenced by a combination of genetic variants with different magnitude effects and treatment-specific additive effects.

**Figure S13.** Enrichment of GO categories for genes with eQTL in Head-Copper, Gut-Copper, or both datasets.

**Figure S14.** Correlations of estimated founder haplotype effects at peak positions of eQTL detected in both the Head-Copper and Gut-Copper datasets skewed positive for the majority of eQTL. A. Estimated founder haplotype effect correlations at eQTL associated with genes with functions unrelated to metal response based on FlyBase annotations were generally continuous and ranged from a small number of strong negative correlations to strong positive correlations. B. Estimated founder haplotype effect correlations at eQTL associated with genes with functions that are related to metal response (oxidative stress response, response to copper, response to xenobiotics, detoxification) generally fell into two groups with a small number of strong negative correlations and a larger number of strong positive correlations.

**Figure S15.** Gene ontology enrichment for potential genotype by environment eQTL that were detected in both control and copper conditions. A. Genes with eQTL that were detected in head tissue under copper and control conditions were enriched for categories related to metal ion transport and stress. B. Genes with eQTL that were detected in gut tissue under copper and control conditions were also enriched for metal-related processes. The size of plotted circles represents the relative number of genes per category. Z-score represents the relative proportion of genes with normalized expression higher or lower than the average (0) under copper conditions.

**Figure S16.** Gene ontology enrichment for potential genotype by environment eQTL. A. Genes with eQTL that were detected in head tissue only under copper conditions were enriched for categories related to metal ion transport and stress. B. Genes with eQTL that were detected in head tissue only under control conditions were enriched for categories related to cellular processes, response to stimulus, and biological regulation. The size of plotted circles represents the relative number of genes per category. Z-score represents the relative proportion of genes with normalized expression higher or lower than the average (0).

**Figure S17.** Gene ontology enrichment for potential genotype by environment eQTL. A. Genes with eQTL that were detected in gut tissue only under copper conditions were enriched for categories related to lipid metabolism. B. Genes with eQTL that were detected in gut tissue only under control conditions were enriched for categories related localization and transport. The size of plotted circles represents the relative number of genes per category. Z-score represents the relative proportion of genes with normalized expression higher or lower than the average (0).

**Figure S18.** Pairwise correlations of gene expression or expression response of *trans* eQTL genes detected in the three Head datasets. The bottom lollipop plot summarizes mean absolute Pearson correlations for each set of *trans* eQTL genes.

**Figure S19.** Pairwise correlations of gene expression or expression response of *trans* eQTL genes detected in the three Gut datasets. The bottom lollipop plot summarizes mean absolute Pearson correlations for each set of *trans* eQTL genes.

**Figure S20.** Loadings for gene included in each *trans* eQTL hotspot that were highly correlated loaded evenly on PC1, which was used as a composite variable in assessment of potential mediators of *trans* eQTL hotspots.

**Figure S21.** Mediation analysis of Head hotspots implicated between zero and three potential candidate genes that may influence the expression or response of the *trans* eQTL genes corresponding to most hotspots. In all QTL maps above, the peak associated with variation in the composite variable representing the *trans* eQTL genes is shown in black. Genes that fell within hotspot intervals are highlighted in green; genes with partial or complete mediating effects on the composite QTL are shown in purple. In the regression plots, estimated haplotype effects for the composite peak and for the *cis* eQTL associated with the potential candidate genes are shown. The purple lines represent the best fit based on a linear model with 95% confidence intervals shaded in grey; statistics provide Pearson correlations and p values are presented without multiple test correction. The DSPR has eight founder strains. Haplotype estimates based on low representation of a founder strain at a given position were not included in the correlation analyses.

**Figure S22.** Mediation analysis of Gut hotspots implicated between one and five potential candidate genes that may influence the expression or response of the *trans* eQTL genes corresponding to most hotspots. In all QTL maps above, the peak associated with variation in the composite variable representing the *trans* eQTL genes is shown in black. Genes that fell within hotspot intervals are highlighted in green; genes with partial or complete mediating effects on the composite QTL are shown in purple. In the regression plots, estimated haplotype effects for the composite peak and for the *cis* eQTL associated with the potential candidate genes are shown. The purple lines represent the best fit based on a linear model with 95% confidence intervals shaded in grey; statistics provide Pearson correlations and p values are presented without multiple test correction. The DSPR has eight founder strains. Haplotype estimates based on low representation of a founder strain at a given position were not included in the correlation analyses.

## Supplemental Tables

**Table S1.** Samples from head and gut tissue were arrayed across four 96-well plates. Sample names are written as “AB_00000_CC” where “A” is tissue (H = head, G = gut), “B” is the level of resistance exhibited by the RIL in Everman et al. (2021) (R = resistant, S = susceptible), “00000” is the five digit RIL ID and “CC” is the treatment (Cu = exposed to copper sulfate, NA = water control). Plates 1 and 2 contained only head samples; plates 3 and 4 contained only gut samples. Copper and control-treated RIL pairs were kept together on tissue-specific plates. Each plate had an even representation of resistant and sensitive RILs. Copper samples (Cu) are shaded in blue; control samples (NA) are not shaded. Black text indicates resistant RILs; red text indicates sensitive RILs. Samples for which library prep failed are indicated with strikethrough text (HR_21180_Cu, HR_21180_NA, GR_21180_Cu, GR_21232_NA, GS_21281_NA).

**Table S2.** Principal components used in correction of datasets in preparation for eQTL mapping.

## Notes

### Competing Interest Statement

The authors have declared no competing interest.

### Summary of Updates

Addition of supplementary figures

